# Deficiency of MicroRNA-181a Results in Transcriptome-Wide Cell Specific Changes in the Kidney that Lead to Elevated Blood Pressure and Salt Sensitivity

**DOI:** 10.1101/2021.03.18.436092

**Authors:** Madeleine R. Paterson, Kristy L. Jackson, Malathi I. Dona, Gabriella E. Farrugia, Bruna Visniauskas, Anna M. D. Watson, Chad Johnson, Minolfa C. Prieto, Roger G. Evans, Fadi Charchar, Alexander R. Pinto, Francine Z. Marques, Geoffrey A. Head

**Affiliations:** Hypertension Research Laboratory, School of Biological Sciences, Faculty of Science, Monash University, Melbourne, Australia; Monash University, Melbourne, Australia; Neuropharmacology Laboratory, Baker Heart and Diabetes Institute, Melbourne, Australia; Drug Discovery Biology, Faculty of Pharmacy and Pharmaceutical Sciences, Monash University Parkville, Australia; Cardiac Cellular Systems Laboratory, Baker Heart and Diabetes Institute, Melbourne, Australia; Department of Physiology, School of Medicine, Tulane University, New Orleans, the USA; Department of Diabetes, Central Clinical School, Monash University, Melbourne, Australia; Monash Micro Imaging, Monash University, Melbourne, Australia; Cardiovascular Disease Program, Biomedicine Discovery Institute and Department of Physiology, Monash University, Melbourne, Australia; Health Innovation and Transformation Centre, Federation University, Ballarat, Australia; Department of Physiology, University of Melbourne, Melbourne, Australia; Centre for Cardiovascular Biology and Disease Research, La Trobe University, Melbourne, Australia; Heart Failure Research Group, Baker Heart and Diabetes Institute, Melbourne, Australia; Department of Pharmacology, Monash University, Melbourne, Australia

**Keywords:** blood pressure, renin, microRNA, single-cell RNA-sequencing, salt, sodium

## Abstract

MicroRNA miR-181a is down-regulated in the kidneys of hypertensive patients and hypertensive mice. *In vitro*, miR-181a is a posttranslational inhibitor of renin expression, but pleiotropic mechanisms by which miR-181a may influence blood pressure (BP) are unknown. Here we determined whether deletion of miR-181a/b-1 *in vivo* changes BP and the molecular mechanisms involved at the single-cell level. We developed a knockout mouse model lacking miR-181a/b-1 genes using CRISPR/Cas9 technology. Radio-telemetry probes were implanted in twelve-week-old C57BL/6J wild-type and miR-181a/b-1 knockout mice. Systolic and diastolic BP were 4-5mmHg higher in knockout compared with wild-type mice over 24-hours (*P<0*.*01*). Compared with wild-type mice, renal renin was higher in the juxtaglomerular cells of knockout mice. BP was similar in wild-type mice on a high (3.1%) versus low (0.3%) sodium diet (+0.4±0.8mmHg) but knockout mice showed salt sensitivity (+3.3±0.8mmHg, *P<0*.*001*). Since microRNAs can target several mRNAs simultaneously, we performed single-nuclei RNA-sequencing in 6,699 renal cells. We identified 12 distinct types of renal cells, all of which had genes that were dysregulated. This included genes involved in renal fibrosis and inflammation such as *Stat4, Col4a1, Cd81, Flt3l, Cxcl16, Smad4*. We observed up-regulation of pathways related to the immune system, inflammatory response, reactive oxygen species and nerve development, consistent with higher tyrosine hydroxylase. In conclusion, downregulation of the miR-181a gene led to increased BP and salt sensitivity in mice. This is likely due to an increase in renin expression in juxtaglomerular cells, as well as microRNA-driven pleiotropic effects impacting renal pathways associated with hypertension.

## Introduction

MicroRNAs (miRs) are small non-coding RNAs known for their ability to regulate entire physiological pathways at the post-transcriptional level, usually by binding to the 3’ untranslated region (UTR) of specific messenger RNA (mRNA) molecules.^1^ MiRs have emerged in clinical medicine as therapeutic drugs and also as biomarkers of disease pathology.^2^ In the cardiovascular field, an important role for miRs has been suggested for hypertension^3^, heart failure, arrhythmias, and acute coronary syndrome.^4^ Of particular interest is miR-181a, which can bind to the 3’UTR of renin mRNA and downregulate renin levels *in vitro*.^5^ MiR-181a is also downregulated in the kidneys of patients with hypertension compared with those with normal blood pressure (BP),^5^ and both circulating and renal miR- 181a are correlated with levels of BP.^6^

Similar to humans, the kidneys of genetically hypertensive mice have lower levels of miR-181a and higher levels of renin compared with their normotensive counterparts.^7^ Moreover, in the human kidney, renin and miR-181a are co-localised in the epithelial cells of the collecting ducts.^6^ Contrary to juxtaglomerular renin, intratubular renin is not released into the circulation. Rather, it has been shown to act in an autocrine or paracrine manner,^8^ regulating the intra-renal renin-angiotensin system (RAS). Furthermore, hyperactivity of the intratubular RAS can affect both sodium sensitivity and BP. However, due to the pleiotropic capacity of miRNAs, it is likely that miR-181a also regulates BP via genes other than renin in a tissue as heterogeneous as the kidney. Thus, understanding the precise pathophysiological and molecular roles of miR-181a may provide a novel modality for preventing cardiovascular disease by treating hypertension.

In the current study, we deleted miR-181a/b-1 to determine its effects on BP and sensitivity to salt. To determine miR-181a mechanism of action we investigated the renal transcriptome at the single-cell level in mice. We describe the development of a new CRIPSR/Cas9 miR-181a/b-1 global knockout (KO) model, which has lower but not total absence of miR-181a, to replicate the levels of miR-181a observed in human hypertension.^5^ We combined physiological and molecular studies in the first report to describe the pleiotropic role of a miR at the single-cell level to control BP.

## Methods

Expanded methodology is available in the online-only supplement.

### Animals

All experiments and surgical procedures were approved by the Alfred Medical Research Education Precinct Animal Ethics Committee (E/1617/2016/B) and conducted in accordance with the Australian Code for the Care and Use of Animals for Scientific Purposes, in line with international standards. Experiments were conducted on age-matched C57BL/6J (wild-type, WT, n=31) and homozygous KO (n=33) mice with global deletion of the gene miR- 181a/b-1 (but miR-181a/b-2 gene was left intact). Food and water were accessible *ad libitum* (Chow pellets, “Irradiated Rat and Mouse Cubes” from Specialty Feeds, Glen Forrest, 0.2% sodium).

### Salt sensitivity

To determine salt sensitivity, a separate group of WT and miR-181a/b-1 KO mice were fed *ad libitum* a low (0.03% sodium) and high (3.0% sodium) sodium diets (Specialty feeds, Glen Forrest). Animals were randomly allocated to receive each of the diets in turn for one week.

### Cardiovascular Measurements

Radiotelemetry transmitters (model TA11PA-C10; Data Sciences International, St Paul, MN) were implanted in 12-week old mice, as detailed in the online-only Data Supplement and per our previous studies (n=12 WT, n=15 KO).^9^ Systolic (SAP), diastolic (DAP) and mean arterial blood pressure (MAP), heart rate (HR) and locomotor activity were measured in conscious, unrestrained mice from 10-days after surgery for a period of 48-hours. To further determine the contribution of the RAS versus the sympathetic nervous system (SNS), we also measured cardiovascular parameters for 30-minutes following administration of pentolinium (5mg/kg; Sigma-Aldrich) to mice pre-treated with enalaprilat (an angiotensin converting enzyme, ACE, inhibitor, at 1mg/kg; Merck & Co.) during the active (dark) and inactive (light) period as previously described^.7^

### Tyrosine hydroxylase staining

Kidneys were fixed with 10% neutral buffered formalin, dehydrated and embedded in paraffin. Tyrosine hydroxylase (TH) staining was performed on kidney sections from WT (n=6) and miR-181a/b-1 KO (n=5) mice. The percentage of TH staining in the cortical tubules was semi-quantitatively assessed as previously described^7^ (see online-only Data Supplement).

### Measurement and localisation of renin mRNA and protein in the kidney

Localisation of renin mRNA was determined in coronal tissue sections using the RNAScope 2.5 Brown Assay (Advanced Cell Diagnostics, Hayward, CA) according to manufacturer’s instructions. Localisation of renin protein was determined using immunostaining by the peroxidase technique. Briefly, formalin fixed mouse kidney sections (2-3 µm) were processed for renin specific immunolocalization as described previously.^10^ Renin kidney immunolocalization was assessed using a polyclonal rabbit renin antibody (provided as a gift by Dr. R. Ariel Gomez, University of Virginia, VA-USA) at a 1:200 dilution. Peroxidase activity was visualized with 0.1% 3,3’-diaminobenzidine tetrahydrochloride (DAB, Sigma, St. Louis, MO), followed by counterstaining and mounting with using aqueous mounting media (Biomeda, Fisher Scientific). Sections were imaged using bright field microscopy (Olympus BX53 Light Microscope, Japan). A macro (developed by C.J.) was used to identify the amount of renin and renin positive juxtaglomerular cells in the FIJI processing package. All juxtaglomerular apparatus (JGA) visible in the slide were counted (average 185 in WT and 192 in KO mice for renin protein and 163 in WT and 177 in KO for renin RNA). However, due to technical difficulties and laboratory/border closures the sample size was small (n=2-4 per group), but number of medulla images assessed per sample were high (average 8.8 regions/sample).

### Single-nuclei RNA-sequencing

Single-cell suspensions were prepared from the kidneys of male WT (n=4) and miR-181a/b-1 KO (n=4) mice. Mice were euthanised, perfused with PBS and the kidneys isolated. Kidneys were weighed and 0.25g was minced. The same region was selected for consistency, and represented a combination of medulla and cortex. The minced kidney tissue was digested with dispase II and collagenase IV, filtered and resuspended. Antibody staining was performed according to the online-only Data Supplement, and live nucleated cells were sorted using fluorescence-activated cell sorting (FACS). Single-nuclei mRNA libraries were prepared per strain using the 10X Genomics pipeline. The Cell Ranger software version 3.1.0 (10X Genomics) was used to process the raw sequencing data before subsequent analyses. Downstream analysis on kidney single-nuclei RNA-sequencing (scRNA-seq) dataset was performed using Seurat R package 3.2.0.^11^ In order to explore the transcriptional heterogeneity and to undertake cell clustering, dimensionality was reduced using principal component analysis (PCA), selecting 40 PC. PC loadings were used as inputs for a graph-based approach to clustering of cells at a resolution of 1.2, and for t-distributed stochastic neighbour embedding (t-SNE) for two-dimensional visualization purposes. Identified clusters were manually annotated based on features corresponding to canonical cell-type genes before merging clusters corresponding to the same cell type (Table S1). Differentially expressed (DE) genes were identified, between WT and KO samples, by identifying the genes expressed in at least 10% of cells in at least one of the groups being compared. To test for differential expression, MAST was used including cellular detection rate as a covariate (MASTcpmDetRate).

### Statistical Analysis

All cardiovascular data were analysed using a mixed model, repeated measure (split-plot) analysis of variance (ANOVA) using Microsoft Excel 2016. The main effect of strain was further analysed using a series of non-orthogonal contrasts. A Bonferroni correction was used to correct for multiple comparisons with a Greenhouse-Geisser correction to adjust for violation of sphericity. GraphPad Prism (version 7) package was used for the statistical analyses and graphing of tissue weight, TH, miR-181a and renin expression. Two-tailed unpaired t-tests were used to compare single values from WT and KO mice. All data are presented as mean ± SEM. Two-tailed *P*<0.05 was considered statistically significant.

## Results

### Deletion of miR-181a increased blood pressure

The successful deletion of the miR-181a/b-1 gene was achieved using CRISPR-Cas9 genome editing (Figure 1). Mice were genotyped using genomic DNA which was confirmed by PCR. miRNA-181a abundance was 10-fold lower in miR-181a/b-1 KO mice compared to WT mice, while heterozygous mice had intermediate levels of miR-181a (*P<0*.*001*, Figure 1). Mice then underwent a 5-week experimental protocol (Figure 1). On average miR-181a/b-1 KO mice had consistently elevated BP over a 24-hour period (*P<0*.*001*, Figure 2). Average 24-hour mean arterial pressure (MAP) was 4.6mmHg higher in miR-181a/b-1 KO mice compared with WT mice (*P<0*.*01*, Figure 2). Similarly, miR-181a/b-1 KO mice had higher 24-hour SAP (mean difference 4.9mmHg, *P<0*.*01*) and DAP (mean difference 4.4mmHg, *P<0*.*01*) than WT mice (Figure 2). The difference in BP between the two strains was most prominent during the dark (active) period, when microRNA-181a/b-1 KO mice (n=15) had greater MAP (+ 5.6 mmHg, *P=0*.*015*), SAP (+6.0 mmHg, *P=0*.*01*), and DAP (+5.4 mmHg, *P=0*.*02*) than WT mice (n=12) (Figure 2). The MAP, SAP and DAP of KO mice were not different to WT during the light inactive period (P>0.05, Figure 2). MicroRNA-181a/b-1 KO mice had a higher heart rate (HR) than WT mice in the dark (active) period (*P=0*.*05*). Activity was 33% greater in miR-181a/b-1 KO mice than WT mice during the dark (active) period (+0.5, *P=0*.*015*, Figure 2). However, no detectable difference in activity was observed during the light (inactive) period or on average over 24-hours. The depressor response to ACE inhibition (enalaprilat) was similar in the two strains (Figure S1). Further, the depressor response to ganglion blockade by pentolinium, after pre-treatment with enalaprilat, was also similar in WT and KO mice during the dark (active) and light (inactive) periods (Figure S1). Deletion of miR-181a did not significantly affect responses to dirty cage, feeding or restraint stress challenges (Figure S1).

**Figure 1.**
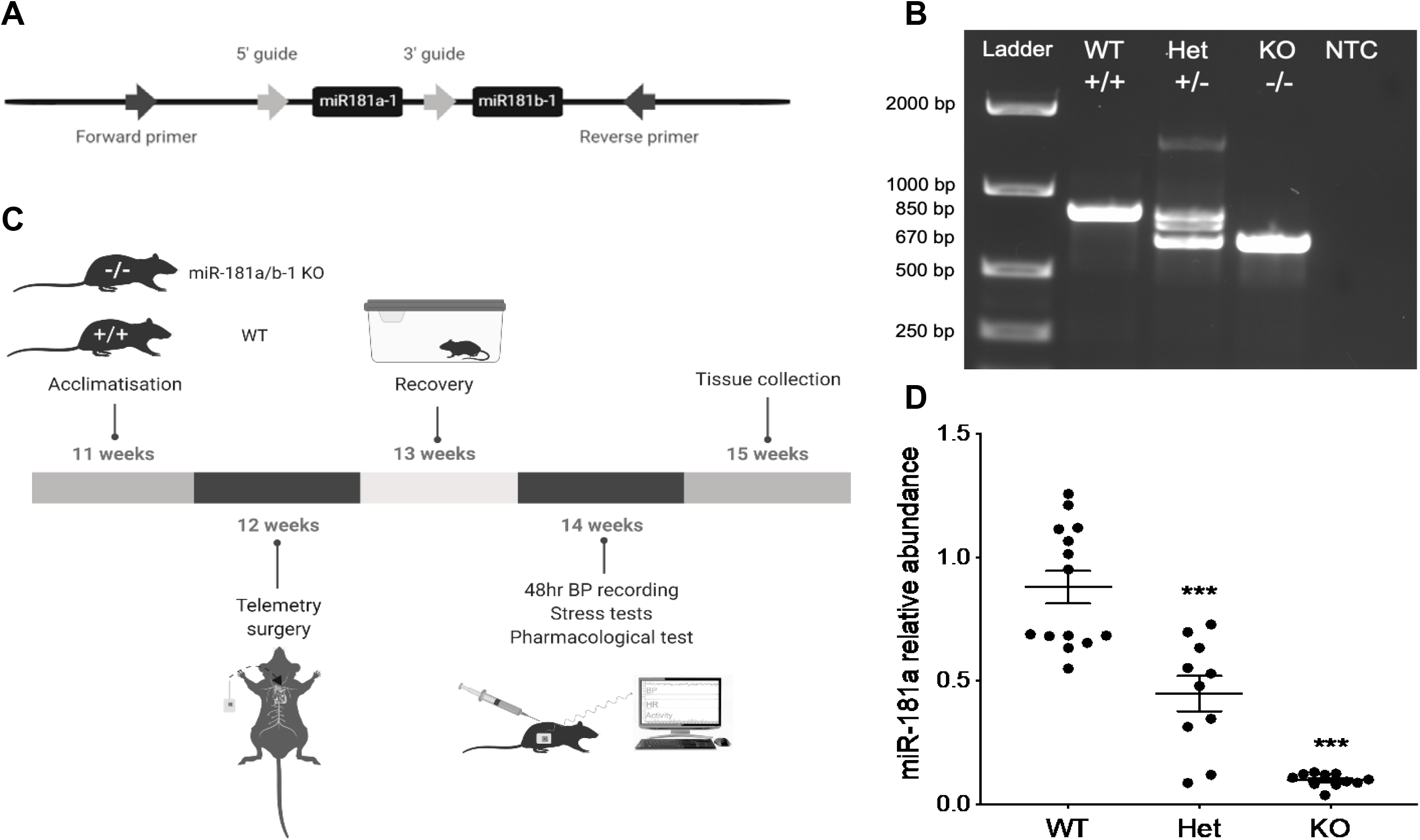
Development and validation of miR-181a knockout. **A**, Example of genotyping strategy used to isolate the miR-181a-1 and miR-181b-1 loci on the wild type allele. **B**, Example of an agarose electrophoresis gel used to identify wildtype (WT), heterozygous (Het) and homozygous (KO) miR-181a/b-1 KO mice, including a non-template control (NTC). Genotypes were distinguished based on the presence of 850 base pair (bp) bands (WT) and 670 bp bands (KO). **C**, Schematic representation of the 5-week experimental protocol timeline. **D**, MicroRNA-181a relative abundance in WT (n=14), miR-181a/b-1 heterozygous (Het, n=10) and homozygous (KO, n=11) KO mice during the active period. Bars represent average values±SEM. Statistical analysis was conducted using one-way analysis of variance. Comparisons are between strains \*\*\**P*<0.001 compared to WT.

**Figure 2.**
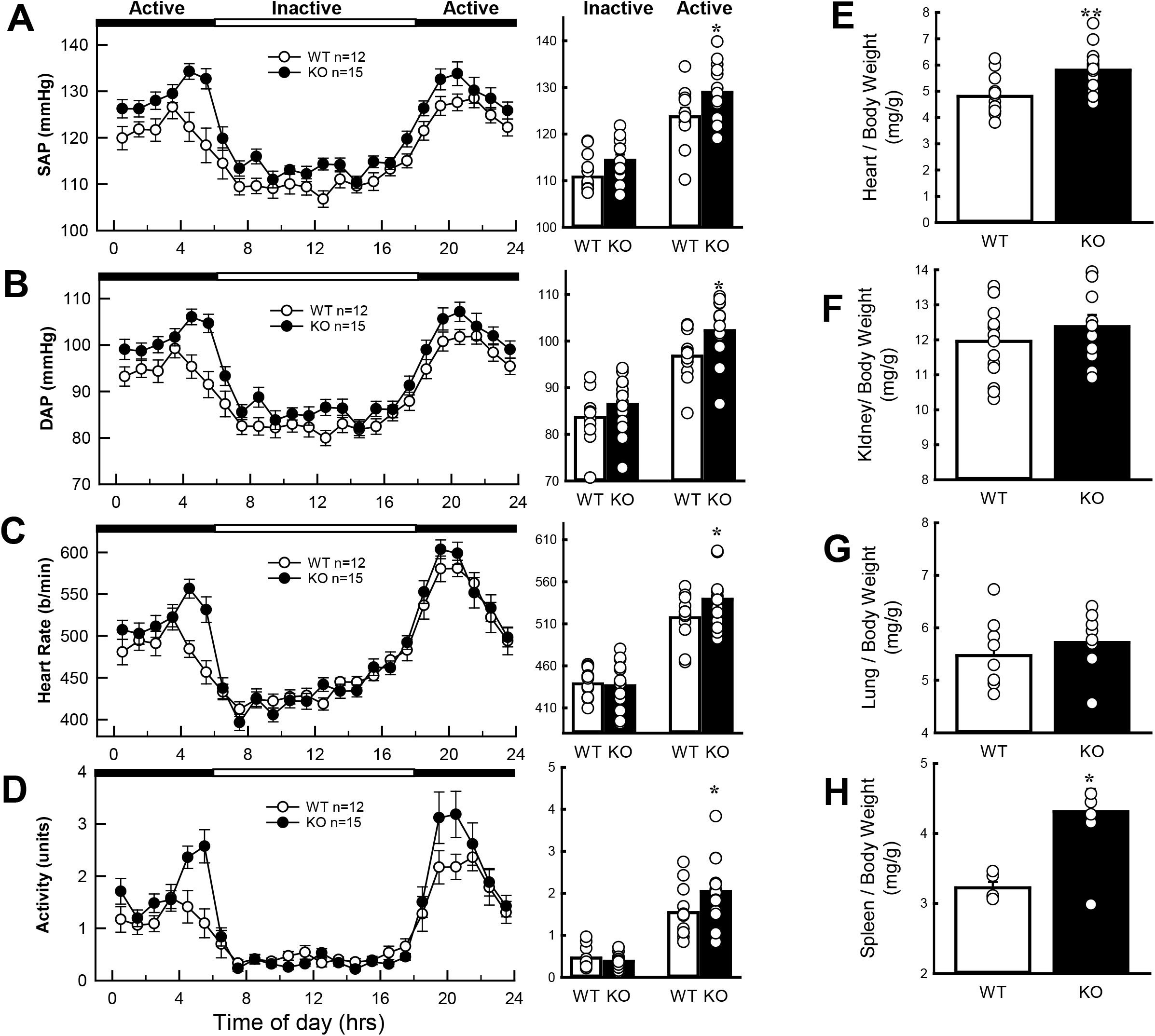
Blood pressure of miR-181a/b-1 knockout. Average 24-hour **A**, systolic blood pressure (SAP), **B**, diastolic blood pressure (DAP), **C**, heart rate (HR) and **D**, activity in WT and miR-181a/b-1 KO mice. The dotted lines show the transition from the active (dark, black panels) periods to the inactive (light, white panel) periods. Histograms (right) represent the average SAP, DAP, HR and activity during the inactive and active periods in each mouse strain. **E**, Heart to body weight ratio (mg/g), **F**, Kidney to body weight ratio (mg/g), **G**, Lung to body weight ratio (mg/g), **H**, Spleen to body weight ratio (mg/g). All values are mean ± SEM. **A-D**, statistical analysis was conducted using between groups, split plot analysis of variance with a Bonferroni and Greenhouse Geisser adjustment. **E-H**, 2 factor ANOVA with post hoc Bonferroni adjusted contrasts two-tailed unpaired t-test. \**P* < 0.05, \*\**P* < 0.01, \*\*\**P* < 0.001 compared to WT.

Food and water intake did not differ between WT and miR-181a/b-1 KO mice (Figure S2). Cardiac weight index, determined as heart to body weight ratio, was 21% greater in miR- 181a/b-1 KO mice than WT mice (*P<0*.*001*, Figure 2). There were no detectable differences in (body-weight adjusted) kidney or lung weight index between miR-181a/b-1 KO and WT mice, while KO mice had larger spleens (+34%, *P*<0.05, Figure 2). TH staining in the kidney, which has been used as a marker of sympathetic innervation, was 2.5-fold greater in miR-18a/b-1 KO mice (5.2±1.1%) than WT mice (2.0±0.3%, *P=0*.*03*, Figure 3). Together, our data suggests miR-181a is involved in BP regulation, leads to cardiac hypertrophy, and increases renal sympathetic activity, but is not involved in the pressor responses to aversive stress.

**Figure 3.**
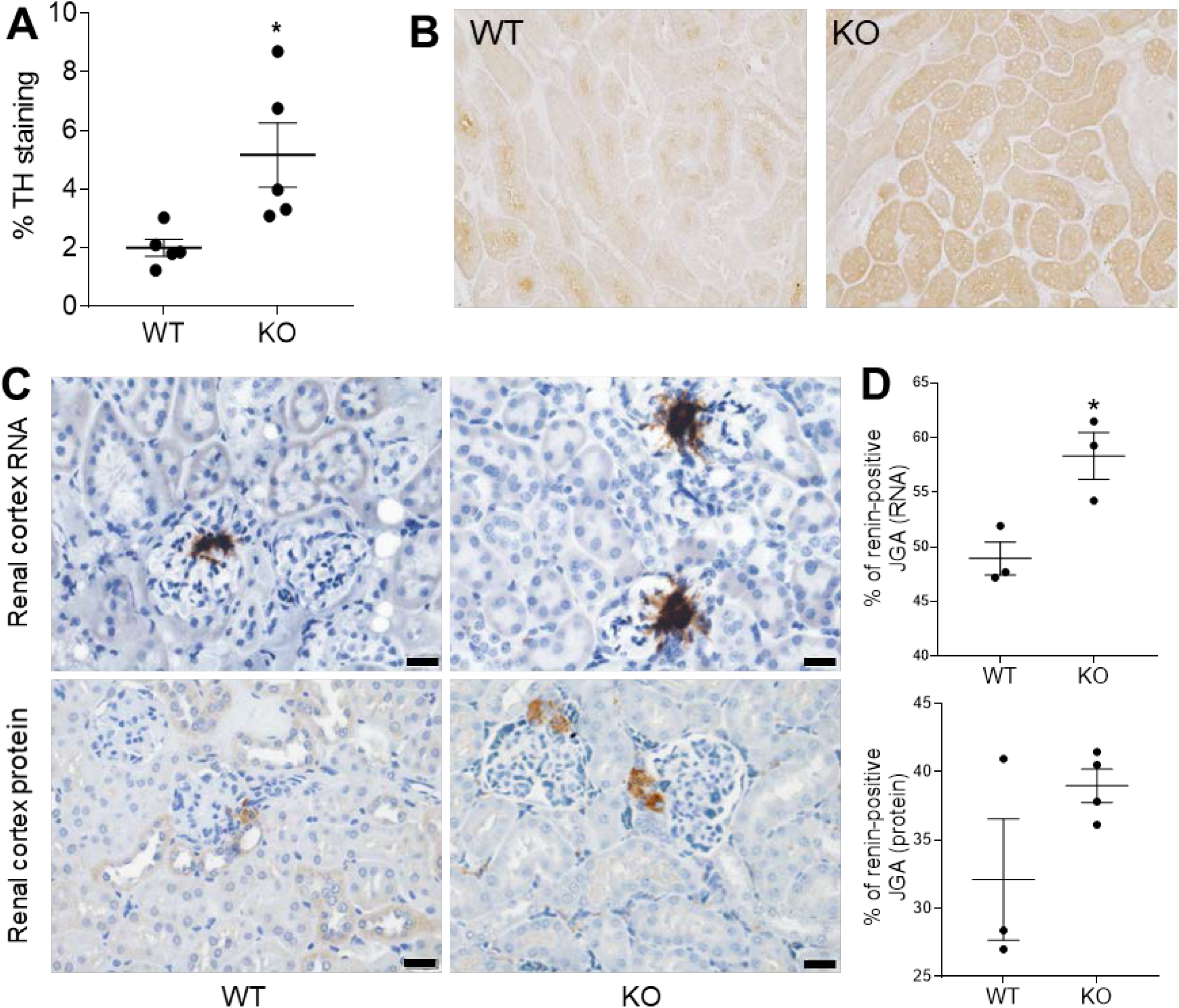
Renal renin and tyrosine hydroxylase in the miR-181a/b-1 knockout. **A**, Percentage area of the image positively stained for tyrosine hydroxylase (TH) in WT and miR- 181a/b-1 KO mice. **B**, Representative micrograph showing TH staining (dark brown) in cortical tubules (scale bar=50 µm) of WT and miR-181a/b-1 KO mice. **C**, Representative images of *in situ* hybridisation (top images) and immunohistochemistry (bottom images), with renin shown as brown stain (scale bars=20µm). **D**, Percentage of renin mRNA positive juxtaglomerular apparatus (JGA) and renin protein in the JGA. Bars represent average values ± SEM. Statistical analysis was conducted using one-way analysis of variance. Comparisons are between strains \**P*<0.05 compared to WT. n=2-5/group.

### Deletion of miR-181a increased renal renin

Since miR-181a binds to and down-regulates renin *in vitro*, it is likely that renin levels are affected. Accordingly, JGA more frequently expressed renin mRNA in miR-181a/b-1 KO mice than WT mice (mean±SEM: WT 48.9±1.5% vs KO 58.3±2.1%, *P=* 0.02, Figure 3) and protein (WT 32.1±4.4% vs KO 39.0±1.2%, *P=*0.14, Figure 3). Higher renin mRNA, but not protein, was also detected in the medulla (WT 4.2±1.0% vs KO 8.7±3.1%, Figure S3). These data suggest that renal renin, particularly in the JGA, is overexpressed in the miR-181a/b-1 KO mice.

### Deletion of miR-181a increased salt sensitivity

WT mice had similar 24-hour MAP when fed a high or low salt diet (mean difference 0.4mmHg, Figure 4). In contrast, miR-181a/b-1 KO mice showed a degree of salt sensitivity (Figure 4). Most notably, DAP was 4.8mmHg greater during the dark (active) period and 3.2mmHg greater during the light (inactive) period in miR-181a/b-1 KO mice fed a high salt diet compared to a low salt diet (*P<0*.*05*, Figure 4). Similarly, a high salt diet led to increased SAP during the dark (active, +3.8mmHg, *P<0*.*05*) but not during the light (inactive, +2.0mmHg, *P=0*.*08*) period, compared with miR-181a/b-1 KO mice fed a low salt diet. There were no detectable differences in urine output, water and food consumption between miR-181a/b-1 KO and WT mice on any of the diets (Figure S2). However, as expected, when fed a high salt diet, both miR-181a/b-1 KO and WT mice had greater water consumption and urine output than when fed a low salt diet (*P<0*.*001*, Figure S2). Together, these results indicate that miR-181a/b-1 KO mice are more salt-sensitive than WT mice.

**Figure 4.**
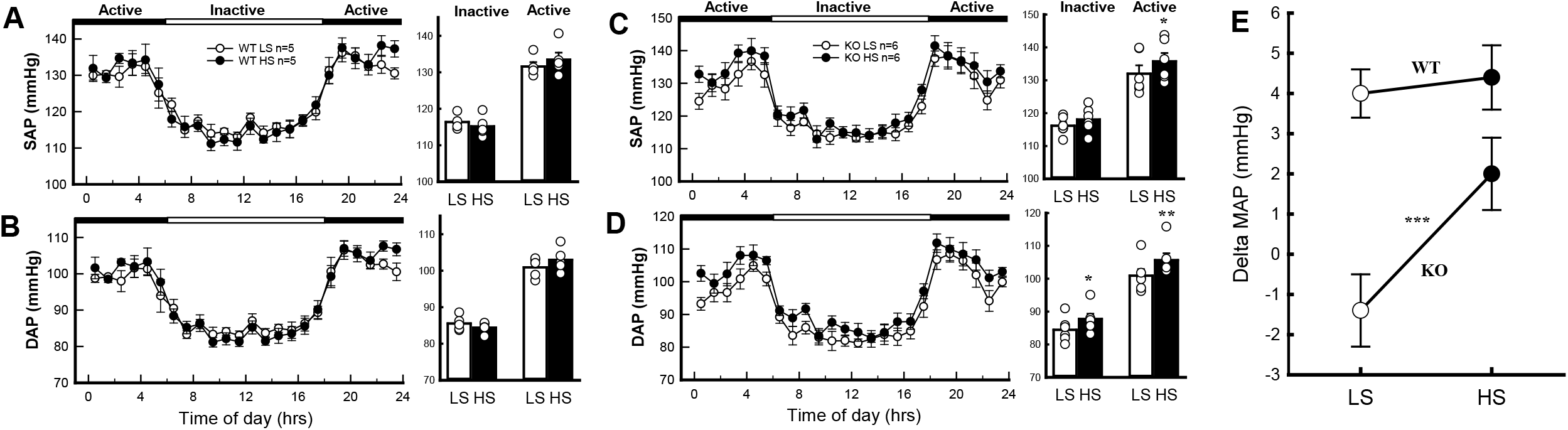
Blood pressure in the miR-181a/b-1 knockout response to salt. Average 24-hour systolic blood pressure (SAP) and diastolic blood pressure (DAP) in WT **A**, and **B**, and miR- 181a/b-1 KO mice **C**, and **D**, fed low (white) and high (black) salt diets. The dotted lines show the transition from the active (dark, black panels) periods to the inactive (light, white panel) periods. Histograms (right) represent the average SAP and DAP during the inactive and active periods in each strain. **E**, Average change in mean arterial pressure (MAP) in WT and miR- 181a/b-1 KO mice in response to low and high salt diets. Statistical analysis was conducted using between groups, split plot analysis of variance with a Bonferroni and Greenhouse Geisser adjustment \**P*<0.05.

### Single-Nuclei RNA-sequencing

To determine the impact of miR-181a KO on renal cell types, we performed scRNA-seq, on nucleated kidney cells (see Online-only Methods) isolated and pooled from four WT and four miR-181a/b-1 KO mice. scRNA-seq yielded 6,699 cells (3,361 cells sequenced at 139,800 reads/cell for WT and 3,338 cells sequenced at 140,517 reads/cell for miR-181a/b-1 KO mice) that passed quality control in our analysis pipeline (Figure 5A and Figure S4A). Following cell clustering, we identified 12 distinct major cell populations in both WT and miR-181a/b-1 KO strains (Figure 5B) including 22 cell populations that corresponded to macrophages, endothelial cells, juxtaglomerular cells, cells of the proximal tubule (PT), dendritic cells (DC), natural killer cells (NK), T cells and B cells (Figure 5C and S4B, Table S1). Importantly, many of the most highly expressed genes in JGA cells were previously identified as part of human reninomas.^12^

**Figure 5.**
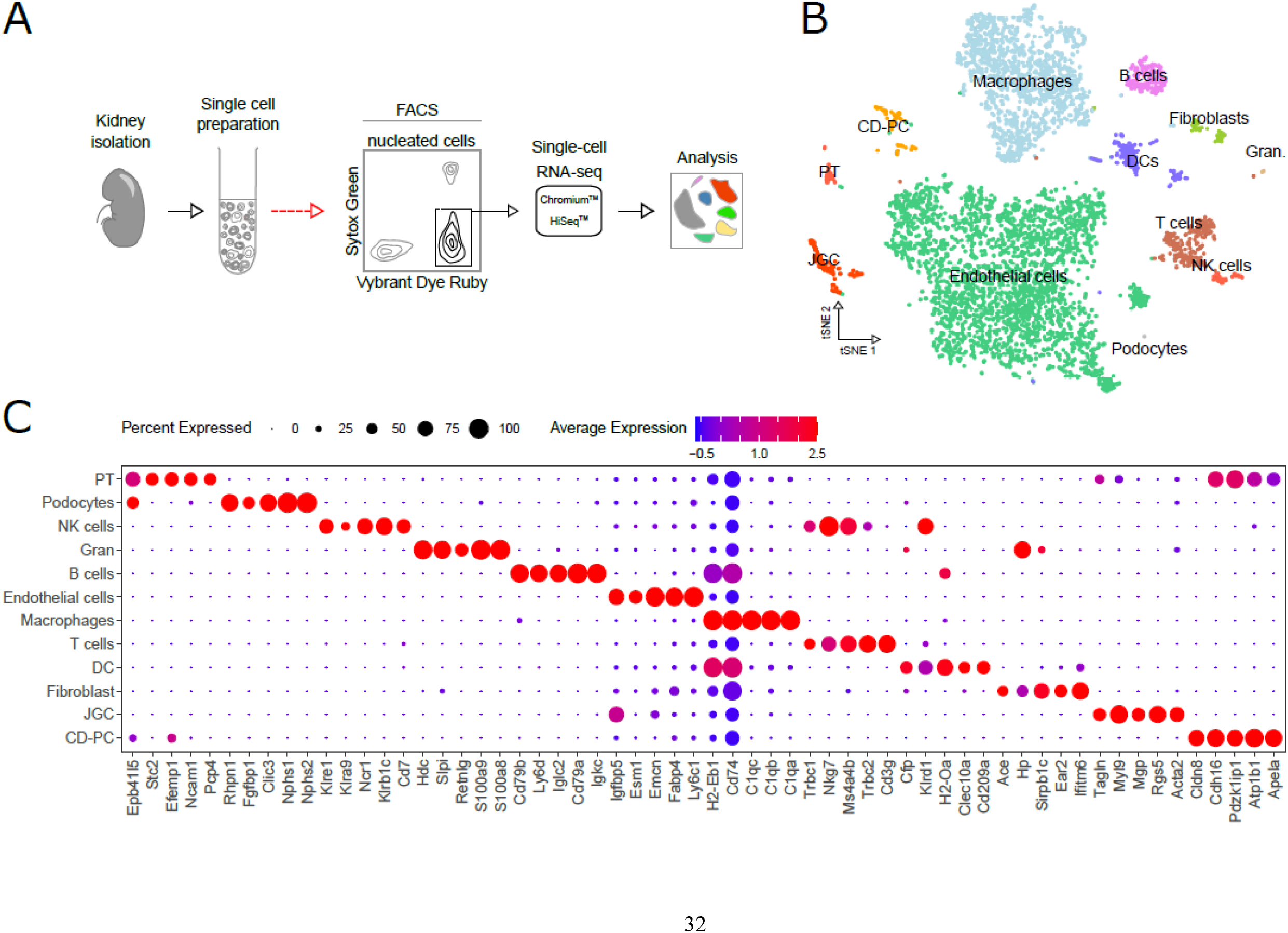
Isolation and analysis of kidney cells by scRNA-seq. **A**, schematic outline of procedure to isolate and analysing cell types from control and miR181a/b-1 KO mouse kidneys. **B**, t-SNE projection of kidney cell types identified by scRNA-seq. **C**, Top-five distinct genes for each cell type using an unsupervised approach (please see Table S1 for top 10 gene markers per cell population). Legend: Gran, granulocytes; PT, proximal tubules; NK cells, natural killer cells; DCs, dendritic cells tubule; JGC, juxtaglomerular cells; CD-PT, collecting duct principal cells.

Analysis of gene expression profiles revealed miR-181a/b-1 KO impacts multiple cell populations to varying degrees with macrophages and endothelial cells most impacted (Figure 6A and Table S2). However, it should be noted that sensitivity for discovering differentially expressed genes is reduced for less abundant cell types in our dataset (Figure S5), which is a common phenomenon in scRNA-seq.^13^ Examination of the top ten up- and down-regulated genes in miR-181a/b-1 KO mice showed that most genes are cell-specifically regulated (Figure 6 and Table S2). Amongst the top up-regulated genes were *Stat4* in B cells (fold change=11, expression validated by real-time PCR in Figure S6) which is associated with inflammation,^14^ collagen genes associated with fibrosis such as *Col4a1* (fold change=1.5) in endothelial cells, *Cd81* a marker of renal-resident macrophages^15^ (fold change=1.2), *Flt3l* found in natural killer cells (only expressed in miR-181a/b-1 KO mice) which exacerbates accumulation of T cells in the kidney,^16^ *Malat1* found in macrophages (fold change=1.2) shown to be higher in hypertensive patients and blockade of *Malat1* was able to lower BP in an animal model,^17^ *Notch2* in proximal tubules (only expressed in miR- 181a/b-1 KO mice) which is involved in nephron development,^18^ and *Polr2f* found in proximal tubules (fold change=2.2) previously associated with established hypertension in a meta-analysis.^19^ Several of the up-regulated genes have been previously demonstrated to have a role in renal injury, inflammation and fibrosis including *Cxcl16*^*20*^ found in distal convoluted tubule cells (fold change=2.1), and *Smad4* found in collecting duct principal cells (fold change=1.3).^21^ The function of most of the down-regulated genes we identified is unknown at least in the context of hypertension and renal function. This is not surprising considering that when miR-181a is downregulated, genes that are directly controlled by this miR would be expected to be upregulated. We still identified a few relevant genes including *Ormdl1* in fibroblasts (absent in KO) associated with BP in a genome-wide association study,^22^ and *Tmem27* in proximal tubule cells (fold change=-1.5) which is an ACE2 analogue also involved in nitric oxide synthesis.^23^

**Figure 6.**
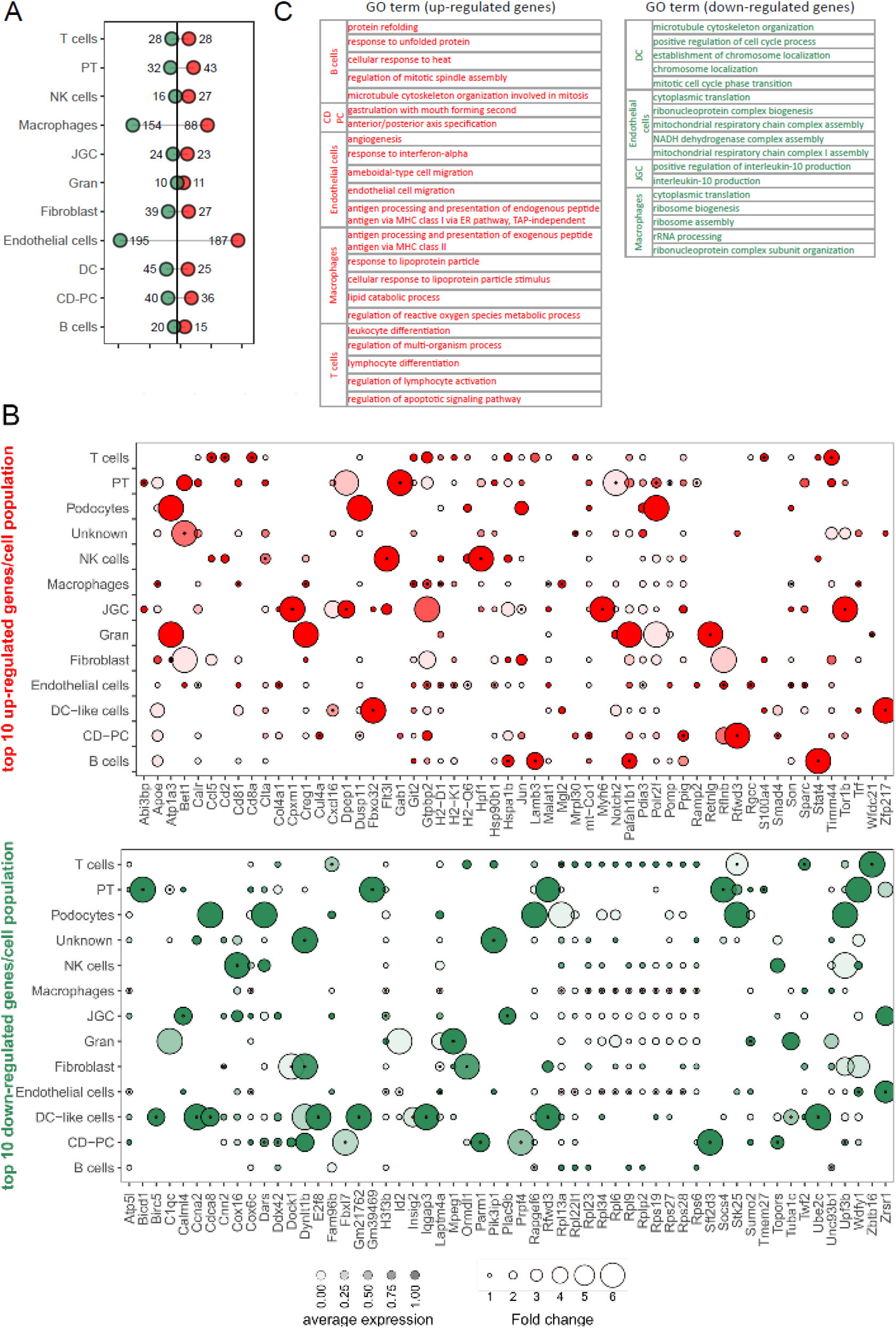
Gene expression changes in miR181a/b-1 KO kidney cell populations. **A**, Lollipop plot summarizing the number of up- and down-regulated genes (uncorrected *P*<0.01, see Table S2). **B**, dot plot summarizing top-ten genes up- or down-regulated within kidney cell types of miR181a/b-1 KO mice relative to control mice. Circle size indicates fold-change; circle saturation indicates relative expression level (dark= high, clear = low); black dots at the centre of circles indicates uncorrected *P*<0.01. **C**, Gene ontology pathways dysregulated in miR-181a/b-1 KO (please see Tables S3 and S4 for complete gene ontology). Legend: CD-PT, collecting duct principal cells DC, dendritic cells; JGC, juxtaglomerular cells; GO; gene ontology; Gran, granulocytes; NK cells, natural killer; PT, proximal tubules.

Gene ontology analyses were then performed to determine pathways altered by miR- 181a/b-1 deficiency. Most identified pathways were down-regulated in dendritic cells, endothelial cells, macrophages and juxtaglomerular cells (Figure 6D, Tables S3 and S5). In juxtaglomerular cells, we observed the down-regulation of pathways involved in interleukin- 10 (IL10) production, which is anti-inflammatory and limits BP increase.^24^ While many down-regulated pathways identified in the other cell types were involved in ribosomal units and translation, in endothelial cells we observed the up-regulation of pathways related to axonogenesis and nerve development, which is consistent with our findings regarding higher levels of TH, which point to higher sympathetic activity. Genes in pathways associated with the activation of the immune system in macrophages, B and T cell populations, and inflammatory response (e.g., interleukin-6 secretion, T cell-mediated cytotoxicity) were also up-regulated. Moreover, there was an up-regulation of pathways involved in the regulation of reactive oxygen species (ROS) metabolic processes in macrophages and B cells. Overall, this data supports an effect of miR-181a/b-1 on many mechanisms that are associated with hypertension including nerve activation, immune system activation, inflammation and ROS.

## Discussion

Here we show that a 10-fold reduction in miR-181a/b, through the use of a new CRISPR/Cas9 KO model, resulted in higher BP, cardiac hypertrophy and caused a degree of salt sensitivity. Similar to our findings from kidneys obtained from humans with confirmed hypertension,^5^ we were able to reduce (but not completely remove) miR-181a expression in this model, while renal renin mRNA and protein appeared increased, particularly in juxtaglomerular cells. These data support that miR-181a exerts post-transcriptional control over renal renin expression. Our single-nuclei transcriptomic analyses, however, defined a pleiotropic effect of miR-181a in BP regulation by the kidneys that goes beyond renin regulation. Indeed, miR-181a also impacted pro-inflammatory IL6 and anti-inflammatory IL10 pathways, known to impact sodium handling.^25^ Finally, the elevated TH staining observed in the kidneys of miR-181a/b-1 KO mice is consistent with increased nerve development identified by scRNA-seq and may point to a role for the SNS in BP regulation in this model. Together, our results support the hypothesis that deletion of miR-181a leads to elevated BP through multiple classical (RAS, SNS) and emerging (non-coding RNA, pro-inflammatory) mechanistic pathways.

We have previously shown that low levels of renal miR-181a are associated with increased BP in human and pre-clinical models.^7,26,27^ Furthermore, circulating plasma levels of miR-181a were also associated with BP in two independent cohorts.^28^ Likewise, expression of miR-181a in monocytes was found to be negatively correlated with systolic BP in obese patients.^29^ Others have demonstrated that mice lacking miR-181a exhibit cardiac hypertrophy and cardiac dysfunction.^30^ These observations and our current data showing cardiac hypertrophy in miR-181a/b-1 KO mice likely reflect the effects of higher BP, since expression of miR-181a in the heart is extremely low (data not shown). Our findings indicate a role of miR-181a in BP regulation *in vivo*, and expand our knowledge about the mechanisms involved at the single-cell level in the kidney. Here, we show for the first time that global deletion of the miR-181a/b-1 genes *in vivo* results in chronically elevated BP over a 24-hour period and pinpoint several novel mechanisms in the kidney. Thus, we demonstrate that under-expression of miR-181a is not merely a consequence of high BP, but rather a driving factor leading to the development of elevated BP.

In the present study, we found that miR-181a/b-1 KO mice had higher levels of renin in the JGA. In addition to JGA, in the human kidney we previously reported that miR-181a co-localizes with renin in the distal nephron.^28^ Collecting duct renin is thought to act within the intrarenal RAS in an autocrine or paracrine manner rather than systemically.^8,32^ In addition to renin, both ACE and angiotensinogen, components required for the synthesis of angiotensin (Ang) II, are present within the renal tubules. Given that tubular Ang II increases epithelial sodium transport within the collecting duct, it has been suggested that overexpression of renin in this region can promote sodium reabsorption and consequent increases in BP. Indeed, mice which overexpress collecting duct renin showed increased BP and salt sensitivity.^33^ Conversely, mice with specific deletion of the *Ren-1c gene* in the collecting duct, or reduced local renin activity due to cell-type specific deficiency of the prorenin receptor (*Atp6ap2*) gene in collecting duct cells, both elicit attenuated Ang II-induced hypertension and decreased distal sodium reabsorption.^34,35^ This hypothesis is supported by the observation in the present study that a high salt diet leads to increased BP in miR-181a/b-1 KO mice but not in WT mice. Further evidence that renin is acting in an autocrine manner is provided by the observed similar depressor response to the ACE inhibitor enalaprilat in WT and miR-181a/b-1 KO mice fed a control diet (Figure S1). Given that enalaprilat inhibits production of circulating Ang II, the equal decrease in MAP in both wild-type and miR-181a/b-1 KO mice suggests that the increased renin, which appears to be in the kidneys of miR-181a/b-1 KO mice, is not released into the circulation. Therefore, it is likely that medullary renin mRNA is promoting greater sodium retention, thus, increasing BP in miR-181a/b-1 KO mice. This also suggests that the decrease in BP is not only due to miR- 181a actions on renin, but that other targets are important. However, to confirm this hypothesis we would need to measure levels of circulating renin, which was not possible in the current investigation.

Our previous findings indicate there may be a link between the SNS and miR-181a.^7^ Indeed, renal denervation lowered BP and normalised expression of miR-181a and renin in Schlager hypertensive (BPH/2J) mice with neurogenic hypertension, but not in normotensive control mice.^36^ Since BPH/2J mice and miR-181a/b-1 KO mice have lower miR-181a levels, one may expect to see altered sympathetic activity in the KO mice.^7^ However, on a normal-salt diet, the depressor response to ganglionic blockade in miR-181a/b-1 KO mice was similar to that of WT mice, both in the active and inactive period. Consistent with this, there were no detectable differences in the sympathetically mediated pressor responses to stress tests between wild-type and miR-181a/b-1 KO mice. Thus, it would appear that the deletion of the miR-181a gene does not affect regulation of the SNS. On the contrary, TH staining, a marker of sympathetic innervation, was greater in miR-181a/b-1 KO mice than wild-type mice, consistent with our scRNA-seq data. Combined with recent findings showing that renal denervation reduced BP and restored levels of miR-181a in the hypertensive mice,^36^ this may imply that the SNS has a role, either directly or indirectly, in the regulation of miR-181a.

We also identified juxtaglomerular cells characterised by the expression of *Ren1, Myl9, Mgp, Rgs5* and *Acta2*, previously characterised in human reninomas,^12^ that had not been previously characterised by scRNA-seq. However, given that miRNAs can bind to and regulate many mRNA targets, a single miRNA can have pleiotropic effects on many different biological pathways above and beyond the renin gene *Ren1*.^37^ Genes co-expressed with miR- 181a were previously determined using next-generation RNA-sequencing.^6^ Renal miR-181a expression was co-expressed with decreased mRNAs common to mitochondrial respiratory function, and with increased expression of mRNAs common to signalling cascades of immunity and inflammation.^6^ In the past decade it has been well established that the pathogenesis of hypertension is due, in part, to immune system dysregulation.^38^ This is relevant as previously miR-181a has been shown to regulate development of T-cells,^39,40^ consistent with the increased spleen size we observed. Furthermore, miR-181a has various immune targets including toll-like receptor 4,^41^ interleukin-1α, interleukin-1β, IL6 and TNFα.^42,43^ For the first time, we have studied the hypertensive mouse kidney at a single-cell level, to further elucidate the pleiotropic effects of miR-181a, and validated that it affects some of these pathways including IL6 signalling pathways.

We acknowledge our study has some limitations. Due to gene proximity, both miR- 181a-1 and miR-181b-1 genes were deleted in mice. There are, however, two genes that code for mature miR-181a: miR-181a-1 and miR-181a-2.^44^ The global deletion of miR-181a/b-1 in mice has been found to have a greater impact on primary and mature miR-181a levels, as well as immune driven function, than KO of miR-181a/b-2.^45^ Given that complete deficiency of miR-181a/b-1 and miR-181a/b-2 in mice does not appear to be compatible with life,^46^ miR-181a/b-1 is probably the main gene responsible for miR-181a expression. Because low (but not total absence of) expression of miR-181a is observed in the kidneys of human hypertensive patients,^5^ the miR-181a/b-1 genes were selected for deletion in the current project. MiR-181a/b was previously shown to regulate endothelial dysfunction via TGF-β signalling in vascular smooth muscle cells, leading to higher pulse wave velocity and systolic BP.^31^ However, human microarray data revealed that, unlike miR-181a, miR-181b is not differentially expressed in human hypertensive kidneys and appears to not regulate renin expression.^5^ Therefore, it is anticipated that the changes in renin expression observed in the miR-181a/b-1 KO mice can be attributed predominantly to the deletion of the miR-181a-1 gene. We also acknowledge that in this study we only analysed the BP and transcriptome of male mice. However, given that data previously obtained from two large cohorts of human hypertensive patients showed no sex differences in miR-181a expression,^6^ we would not expect to see a difference in our model.

### Perspectives

Our findings demonstrate that deficiency of miR-181a/b via global deletion of miR-181a/b-1 gene leads to elevated BP and some salt sensitivity in mice, and the mechanisms involved at the single-nuclei level in the kidney. Our findings show that a single miRNA can influence a wide array of pathways in several cell types in a tissue as heterogeneous as the kidney, leading to a complex phenotype. While there are currently antihypertensive agents available which target the RAS, understanding how down-regulation of miR-181a can lead to elevated BP and salt sensitivity provides opportunities for the development of novel therapies to prevent, diagnose and treat hypertension.

## Acknowledgments

We acknowledge the technical assistance of John-Luis Moretti. We thank Dr. R. Ariel Gomez from University of Virginia for the anti-renin antibody, and the Australian Genome Research Facility (AGRF) for high throughput sequencing, and the support it receives from the Commonwealth.

