## Supplemental file for "Deficiency of MicroRNA-181a Results in Transcriptome-Wide Cell Specific Changes in the Kidney that Lead to Elevated Blood Pressure and Salt Sensitivity"

**Short title***:* MicroRNA-181a and Blood Pressure

Madeleine R. Paterson, BSc[1,2]; Kristy L. Jackson, PhD[2,3]; Malathi I. Dona, PhD[4]; Gabriella E. Farrugia, BSc[4]; Bruna Visniauskas, PhD[5]; Anna M. D. Watson, PhD[2,6]; Chad Johnson, MSc;[7] Minolfa C. Prieto, MD, PhD[5]; Roger G. Evans, PhD[8]; Fadi Charchar, PhD[9,10]; Alexander R. Pinto, PhD [3,11]; Francine Z. Marques, PhD[1,12*]; Geoffrey A. Head, PhD[2,13*]

[1]Hypertension Research Laboratory, School of Biological Sciences, Faculty of Science, Monash University, Melbourne, Australia; Monash University, Melbourne, Australia; [2]Neuropharmacology Laboratory, Baker Heart and Diabetes Institute, Melbourne, Australia; [3]Drug Discovery Biology, Faculty of Pharmacy and Pharmaceutical Sciences, Monash University Parkville, Australia; [4]Cardiac Cellular Systems Laboratory, Baker Heart and Diabetes Institute, Melbourne, Australia; [5]Department of Physiology, School of Medicine, Tulane University, New Orleans, the USA; [6]Department of Diabetes, Central Clinical School, Monash University, Melbourne, Australia; [7]Monash Micro Imaging, Monash University, Melbourne, Australia; [8]Cardiovascular Disease Program, Biomedicine Discovery Institute and Department of Physiology, Monash University, Melbourne, Australia; [9]Health Innovation and Transformation Centre, Federation University, Ballarat, Australia; [10]Department of Physiology, University of Melbourne, Melbourne, Australia; [11]Centre for Cardiovascular Biology and Disease Research, La Trobe University, Melbourne, Australia; [12]Heart Failure Research Group, Baker Heart and Diabetes Institute, Melbourne, Australia; [13]Department of Pharmacology, Monash University, Melbourne, Australia.

*Contributed equally as senior authors

**Correspondence to:** Geoffrey A. Head, Neuropharmacology Laboratory, Baker Heart and Diabetes Research Institute, P.O. Box 6492, St Kilda Road Central, Melbourne, Victoria 8008, Australia.

### Supplement Materials and Methods

#### *Animals*

#### *Generation of miR-181a/b-1 knock out mice*

MicroRNA-181a/b-1 knockout (KO) mice were generated using CRISPR-Cas9 genome editing at the Australian Phenomics Network, Monash Animal Research Platform, Monash University, Clayton. Guide-RNAs were designed using a web-based CRISPR design tool from the Massachusetts Institute of Technology (www.crispr.mit.edu). The Cas9 RNA, 3' (ACCGCAAAGCAGGACCCGAC AGG) and 5' (TGTGTGACAGGTTTGGTTAA AGG) guide-RNAs were microinjected into one-cell-stage C57Bl/6J mouse zygotes. The manipulated embryos were then transferred into the oviducts of pseudo-pregnant foster mothers and allowed to develop. MicroRNA-181a/b-1 heterozygous offspring were then identified and selected as founders for the miR-181a/b-1 KO breeding colony. Founding heterozygous mice were crossbred to establish wild-type (WT) and miR-181a/b-1 KO homozygous and heterozygous mice lines. Both WT littermates of miR-181a/b-1 KO mice and stock C57/BL6 mice were selected as control animals for the present study.

All mice were co-housed with littermates prior to telemetry surgery, after which they were housed singly to allow for individual measurements of cardiovascular variables and locomotor activity. Mice were housed in a temperature and humidity-controlled facility where the average temperature was 24° C and average humidity was 40%. Mice were exposed to a 12:12 hour light and dark cycle (1am–1pm light/day). Food and water were accessible *ad libitum* (Chow pellets, "Irradiated Rat and Mouse Cubes" from Specialty Feeds, Glen Forrest, 19.0% protein, 5.0% fat, 5.0% fibre, 0.2% sodium) with the exception of fasting periods prior to the acute feeding experiment. All experimental and surgical procedures were approved by AMREP Animal Ethics Committee and were conducted in accordance with the Australian Code of Practice for the Care and Use of Animals for Scientific Purposes.

#### *Genotyping of miR-181a/b-1 knock out mice*

Mice from the miR-181a/b-1 colony were genotyped using genomic DNA from tail biopsies collected at 3 weeks of age. DNA was extracted using the PureLink® Genomic DNA kit (Life Technologies, Carlsbad, CA) and amplified using Taq polymerase (Bioline Pty Ltd, Alexandria, NSW, Australia), 200nM of forward (5' ACATGCGTCCTTGCAGTTCT 3') and reverse (5' CCCCATTTGTAACCCCCAGG 3') primers. The PCR cycle was as follows: 1 cycle at 94 °C for 2 minutes, followed by 35 cycles at 94 °C for 30 seconds, 62 °C for 30 seconds and 72 °C for 1 minute, followed by a final cycle at 72 °C for 5 minutes. The primers amplified a product of approximately 850 base pairs from the WT allele and a product of approximately 670 base pairs from the homozygous KO allele. The same genotyping protocol was used to identify colony founders and the mice subsequently bred in the colony.

#### *Telemetry transmitter implantation*

At 12 weeks of age mice underwent surgery to implant a radio-telemetry probe (model TA11PA-C10; Data Sciences International, St Paul, MN). Prior to surgery the mice were weighed and the analgesic carprofen (5 mg/kg s.c, Rimadyl, Animal Health, West Ryde, NSW, Australia) was administered. Anaesthesia was induced by inhalation of isoflurane (Forthane, Abbott, Botany, NSW, Australia) via an open circuit inhalation box (4% induction and 1.5-2% maintenance). Radio-telemetry probes were implanted in mice using aseptic techniques. Mice received an injection of the local anaesthetic bupivacaine (0.01 mL s.c., 0.5 % w/v) at the site of incision. The ventral neck area was shaved, sterilized and a ventral midline incision was made from the lower mandible posteriorly to the sternum. The left carotid artery was exposed by gently separating the submaxillary glands and any surrounding connective tissue. The carotid artery was then occluded using a silk suture (3/0 silk ties; Dysilk, Dynek, SA, Australia) at the end of the artery distal to the head. The catheter was inserted through a small incision into the lumen of the vessel and passed to a point where the tip lay in the aortic arch. A third suture was used to secure the catheter in place. Tissue glue (3M Vetbond, Data Sciences International, St. Paul, MN, U.S.A.) was used to secure the three sutures in place. A subcutaneous pouch was made through the same ventral neck incision and the body of the probe was inserted along the mouse's right flank. The incision site was closed with three silk sutures. Mice were given a 10-day recovery period prior to the cardiovascular measurements.

***Cardiovascular measurements***

The cardiovascular variables BP, HR and locomotor activity were recorded in conscious, unrestrained mice. Mice equipped with radio-telemeters were singly housed on top of PhysioTel receivers (Model: RLA1020, Data Sciences International, St Paul, USA). The receivers detected the radio signal emitted from the probe and transmitted those data to an analog-digital data acquisition card (8024E, National Instruments). The data were sampled on the data acquisition card at 1000 Hz. Two-second average, beat-to-beat and binary wave-form measurements of BP, HR and locomotor activity were recorded and analysed in programs written in labVIEW. The "Data processing' program was used to calculate the average cardiovascular and locomotor data between specific time periods and over selected time intervals (e.g. average HR in 10-minute intervals over the 60 minutes in which the animal underwent a stress test). The 'Circad' program was used to filter and collate the 24-hour BP data by calculating hourly averages from a 72-hour recording period.

***Stress tests and pharmacological assessments***

After a 10-day recovery period the cardiovascular responses to 60-minute exposure to restraint and dirty-cage switch and the non-aversive feeding test were recorded during the inactive (light) period on separate days.[1] Cardiovascular parameters were measured for 30-minutes following administration of pentolinium (5mg/kg; Sigma-Aldrich) to mice pre-treated with enalaprilat (1mg/kg; Merck & Co.). This test was performed both during the active (dark) and inactive (light) periods.

***Metabolic analysis of mice on a low, normal and high salt diet***

Wild-type (n=6) and miR-181a/b-1 KO (n=6) mice were randomly allocated to each of the two salt diets for a week-long period. At the conclusion of each week mice were housed individually in metabolic cages to allow for the measurement of urine production and food and water consumption. Urine sodium and potassium concentration was quantified using indirect ion-selective electrodes (Alinity, Abbott, Abbot Park, IL).

***Real-time PCR***

RNA was processed as previously described.[2,3] Real-time PCR was performed in a QuantStudio 7 instrument (Thermo Fisher Scientific) using Fast SYBR® Green Master Mix (Thermo Fisher Scientific) and custom-made primers for *Stat4* and housekeeping genes 18S and *Actb* (Table S5). Gene expression was analysed as previously described.[2,3]

***Tyrosine Hydroxylase Immunohistochemical Staining***

Staining for tyrosine hydroxylase (TH) was conducted as described previously.[3,4] 4µm paraffin sections (from 10% neutral buffered formalin fixed kidneys) were dewaxed and rehydrated through ethanol before citrate buffer heart antigen retrieval was performed. Sections were then incubated overnight with rabbit anti-TH (polyclonal, 1:100, Merck-Millipore, Temecula, CA, USA) or normal rabbit IgG (1:400 Santa Cruz, Santa Cruz, CA, USA). Subsequently sections were incubated with biotinylated anti-rabbit (Abacus ALS; Vector Laboratories, Burlingame, CA, USA), avidin-biotin complex (Vector) and visualised with 3,3′-diaminobenzidine tetrahydrochloride/H2O2 (DAB; Sigma-Aldrich, St Louis, MO, USA). The percentage of TH staining in cortical tubules was assessed under identical light/exposure (10 images per animal; Eclipse Ci with DS-Ri2 camera, Nikon, Tokyo, Japan). Sections incubated with normal rabbit IgG showed negligible staining (data not shown). The percentage area of the image that stained positively for TH was assessed in a blinded manner based on a red, green and blue threshold (Image Pro Analyzer 7.0 software; Media Cybernetics, Silver Spring, MD).

***Single Cell Library Preparation***

Animals were euthanized with intraperitoneal administration of pentobarbitone and perfused with PBS. Kidneys were harvested, cross-sectioned separating the anterior and posterior sections, and weighed. Up to 0.25g was minced and subject to enzymatic digestion in 3 ml of 2mg/mL collagenase IV (LS004188, Worthington Biochem) and 1 mg/ml dispase II (04942078001, Roche) in 0.9 mM $CaCl_2$ in PBS. Tissue was incubated at 37°C for a total of 45 min with trituration using wide bore tips at 15 minute points. Post-incubation, cell suspension was filtered through 75 µm mesh into 10 mL PBS in 15 ml conical tube. Cells were cleared of debris by centrifugation at 200×*g*, at 4°C for 15 minutes with the breaks off. Supernatant aspirated, and cell pellet washed with *Fx buffer* (2% FCS (heat inactivated), 0.9 mM $CalCl_2$ in 1XHBSS). Cells were stained as instructed with Sytox™ Green (S34860, Invitrogen) and Vybrant™ DyeCycle™ (V10273, Invitrogen) with the addition of verapamil and FACS sorted for live, nucleated cells. Libraries were prepared from sorted cells according to the 10x Genomics Single cell gene expression 3` v2 kit and sequenced in an Illumina HiSeq instrument.

***Analysis of single-cell RNA-Seq data***

Cell ranger version 3.1.0 (10X Genomics) was used to process raw sequencing data files by converting basecall files to fastq format, aligning sequencing reads to mm10 transcriptome and quantifying the expression of each transcript in each cell. There were 6,699 cells that passed quality control steps implemented in Cell Ranger. Downstream analysis on kidney single-cell RNA sequencing (scRNA-seq) dataset was performed in R version 3.6.0 using Seurat R package version 3.2.0.[5] As a further quality-control measure, cells meeting any of the following criteria: <200 unique genes expressed, >10,000 UMIs, or >10% of reads mapping to mitochondria were filtered out. These steps removed an additional 1169 cells, resulting in a final dataset of 5,530 cells (2,684 and 2,846 cells in WT and KO respectively). The Seurat normalization and scaling based on highly variable genes (n=2,000) were then performed with default parameters. Further, Seurat cell cycle regression was performed based on calculated S.Score and G2M.score to regress out the effect of cell cycling genes in the dataset. In order to explore the transcriptional heterogeneity and to undertake cell clustering, dimensionality reduction was performed using PCA with 40 PCs. PC loadings were used as inputs for a graph-based approach to clustering of cells at a resolution of 1.2, and for t-distributed stochastic neighbour embedding (t-SNE) for two-dimensional visualization purposes. Identified clusters were manually annotated based on features corresponding to canonical cell-type genes.

#### *Differential expression analysis*

In order to identify differential gene expression, genes expressed in at least 10% of cells in at least one of the groups being compared were first filtered. The differential expression (DE) testing method MAST with cellular detection rate as covariate (MASTcpmDetRate) was used to identify DE genes between groups. The uncorrected p value cut-off 0.01 was used to determine statistically significantly and differentially expressed genes.

#### *Gene Ontology analysis*

Gene Ontology (GO) enrichment analysis for differentially expressed gene lists (uncorrected p <0.01) was performed using the enrichGO function from ClusterProfiler R package version 3.12.0. All mappings were based on the data from *org.Mm.eg.db*: Genome wide annotation for Mouse, R package version 3.8.2. The over-representation of GO Biological Process terms (GO-BP) was calculated using the list of genes identified in the experiment as the background gene list for *Mus musculus* with minimum and maximum gene set sizes 10 and 500, respectively. GO-BP terms that have semantic similarity higher than the cut-off of 0.7 were treated as redundant terms and discarded using simplify function from clusterProfiler R package. The Benjamini-Hochberg adjusted p-value cut-off of 0.05 was used to determine statistically significant GO-BP terms.

### Tables

**Table S1.** Top 10 marker genes used for identification of cell types from the single-cell RNA-sequencing data. Full marker table available upon request.

| **Cell type** | **Marker genes** |
|---|---|
| Collecting duct principal cells[6] | *Apela, Atp1b1, Pdzk1ip1, Cdh16, Cldn8, Wfdc2, Plet1, Spint2, Pcbd1, Bicc1* |
| Proximal tubule cells[6] | *Pcp4, Ncam1, Efemp1, Stc2, Epb41l5, Bdh2, Wnt16, Sod3, Osr2, Steap2* |
| Juxtaglomerular cells[7] | *Acta2, Rgs5, Mgp, Myl9, Tagln, Tpm2, Myh11, Ndufa4l2, Mustn1, Fxyd1, Ren1* |
| Podocyte[6] | *Nphs2, Nphs1, Clic3, Fgfbp1, Rhpn1, B3galt2, Robo2, Wipf3, Wt1, Magi2* |
| Fibroblasts[6,8] | *Ifitm6, Ear2, Sirpb1c, Hp, Ace, F13a1, Gm21188, Gm9733, Trem3, Gm5150* |
| Endothelial[8] | *Ly6c1, Fabp4, Emcn, Esm1, Igfbp5, Plpp1, Ly6a, Meis2, Egfl7, Kdr* |
| Macrophages[8] | *C1qa, C1qb, C1qc, Cd74, H2-Eb1, H2-Aa, H2-Ab1, Ctss, Wfdc17, Fcer1g* |
| B cells[8] | *Igkc, Cd79a, Iglc2, Ly6d, Cd79b, Ms4a1, Iglc3, Ighd, H2-DMb2, Iglc1* |
| T cells[8] | *Cd3g, Trbc2, Ms4a4b, Nkg7, Trbc1, Cd3d, Trac, Cxcr6, Hcst, Thy1* |
| Dendritic cells[8] | *Cd209a, Clec10a, H2-Oa, Klrd1, Cfp, Mcemp1, Jaml, Tnni2, Kctd14, Rgs18* |
| Natural killer cells[8] | *Cd7, Klrb1c, Ncr1, Klra9, Klre1, Klra8, Il2rb, Prf1, Klrc1, Klrc2* |
| Granulocytes[8] | *S100a8, S100a9, Retnlg, Slpi, Hdc, Clec4e, Clec4d, Wfdc21, Mmp9, Acod1* |

**Table S2.** List of the top 10 genes differentially expressed across all cell types identified by single-cell RNA-sequencing.

| Gene name | Cell type | P-value | Log2Fold Change | Expression in WT | Expression in KO |
|---|---|---|---|---|---|
| *Rpl13a* | Macrophages | 1.27E-29 | -0.53 | 18.42 | 12.79 |
| *H2-D1* | Endothelial cells | 2.48E-24 | 0.39 | 24.37 | 32.03 |
| *H3f3b* | Endothelial cells | 1.27E-23 | -0.41 | 16.08 | 12.13 |
| *Malat1* | Macrophages | 1.14E-21 | 0.23 | 621.07 | 727.25 |
| *Zrsr1* | Endothelial cells | 1.78E-21 | -2.33 | 0.61 | 0.12 |
| *Rgcc* | Endothelial cells | 3.04E-20 | 0.64 | 4.92 | 7.64 |
| *H2-K1* | Endothelial cells | 9.46E-18 | 0.42 | 12.23 | 16.35 |
| *Rpl22l1* | Endothelial cells | 9.99E-17 | -0.53 | 4.91 | 3.39 |
| *A430106G13Rik* | Endothelial cells | 5.05E-16 | | 0.00 | 0.16 |
| *Rpl13a* | Endothelial cells | 6.38E-16 | -0.39 | 11.20 | 8.55 |
| *Mgl2* | Macrophages | 6.41E-16 | 0.81 | 3.68 | 6.46 |
| *Col4a1* | Endothelial cells | 1.9E-14 | 0.59 | 3.09 | 4.65 |
| *Id2* | Endothelial cells | 6.56E-14 | -0.77 | 3.23 | 1.89 |
| *Rpl23* | Macrophages | 9.17E-13 | -0.24 | 35.35 | 29.91 |
| *Rps27* | Macrophages | 1.24E-12 | -0.24 | 74.15 | 62.57 |
| *Rpl9* | Macrophages | 2.07E-12 | -0.29 | 27.56 | 22.51 |
| *Rps19* | Macrophages | 3.51E-12 | -0.21 | 41.68 | 35.93 |
| *Rpl34* | Macrophages | 1.11E-11 | -0.28 | 31.99 | 26.43 |
| *Rps6* | Macrophages | 1.56E-11 | -0.23 | 27.57 | 23.48 |
| *H3f3b* | Macrophages | 1.76E-10 | -0.36 | 17.01 | 13.23 |
| *Dynlt1b* | Dendritic cells | 3.51E-07 | -3.07 | 0.60 | 0.07 |
| *Pin4* | Dendritic cells | 2.17E-06 | -1.80 | 1.18 | 0.34 |
| *Stat4* | B cells | 0.00002 | 3.46 | 0.11 | 1.24 |
| *Pomp* | Proximal tubule cells | 0.00002 | 0.13 | 2.53 | 2.78 |
| *Ccl5* | T cells | 0.00003 | 1.14 | 44.19 | 97.54 |
| *Bicd1* | Proximal tubule cells | 0.00003 | -4.07 | 0.62 | 0.04 |
| *Upf3b* | Collecting duct principal cells | 0.00003 | -0.39 | 1.35 | 1.03 |
| *Smad4* | Collecting duct principal cells | 0.00004 | 0.38 | 0.27 | 0.35 |
| *Iqgap3* | Dendritic cells | 0.00004 | | 0.15 | 0.00 |
| *Cul4a* | Collecting duct principal cells | 0.00006 | 0.64 | 0.36 | 0.56 |
| *Zbtb16* | T cells | 0.00007 | | 0.33 | 0.00 |
| *Rpl13a* | T cells | 0.0001 | -0.37 | 48.41 | 37.48 |
| *Calml4* | Juxtaglomerular cells | 0.0001 | -2.04 | 1.07 | 0.26 |

| | | | | | |
|---|---|---|---|---|---|
| *Plac9b* | Juxtaglomerular cells | 0.0001 | -1.96 | 2.08 | 0.54 |
| *Pdcl* | Dendritic cells | 0.0001 | 3.18 | 0.03 | 0.29 |
| *Abi3bp* | Proximal tubule cells | 0.0002 | 0.57 | 0.49 | 0.73 |
| *Stk25* | T cells | 0.0002 | -2.35 | 0.57 | 0.11 |
| *Socs4* | Proximal tubule cells | 0.0002 | | 0.45 | 0.00 |
| *Laptm4a* | Fibroblast | 0.0002 | -0.92 | 6.19 | 3.26 |
| *Dock1* | Fibroblast | 0.0003 | | 0.33 | 0.00 |
| *Pafah1b1* | B cells | 0.0003 | 1.83 | 0.48 | 1.70 |
| *Cep295* | Dendritic cells | 0.0003 | -2.81 | 0.43 | 0.06 |
| *Polr2f* | Proximal tubule cells | 0.0003 | 1.14 | 1.15 | 2.53 |
| *Ramp2* | Proximal tubule cells | 0.0003 | 0.85 | 0.37 | 0.67 |
| *Jun* | Juxtaglomerular cells | 0.0003 | 0.95 | 0.52 | 1.00 |
| *Cpxm1* | Juxtaglomerular cells | 0.0003 | | 0.00 | 0.86 |
| *Ormdl1* | Fibroblast | 0.0003 | | 0.35 | 0.00 |
| *Atp1a3* | Fibroblast | 0.0003 | 0.00 | 0.78 | 0.78 |
| *Lamb3* | B cells | 0.0004 | 1.99 | 0.52 | 2.08 |
| *S100a4* | T cells | 0.0004 | 0.81 | 4.86 | 8.49 |
| *Tor1b* | Juxtaglomerular cells | 0.0004 | 3.95 | 0.08 | 1.19 |
| *Stil* | Dendritic cells | 0.0004 | | 0.06 | 0.00 |
| *Hspa1b* | B cells | 0.0004 | 1.45 | 1.67 | 4.56 |
| *Ppig* | Collecting duct principal cells | 0.0004 | 1.30 | 0.73 | 1.81 |
| *Dpep1* | Juxtaglomerular cells | 0.0004 | 2.03 | 0.98 | 4.01 |
| *Clta* | Natural killer cells | 0.0004 | 1.21 | 1.48 | 3.43 |
| *Tmem27* | Proximal tubule cells | 0.0004 | -0.58 | 5.36 | 3.59 |
| *Capn15* | T cells | 0.0004 | | 0.00 | 0.21 |
| *Gab1* | Proximal tubule cells | 0.0004 | | 0.00 | 0.51 |
| *Unc93b1* | B cells | 0.0005 | -0.46 | 3.79 | 2.75 |
| *Rfwd3* | Collecting duct principal cells | 0.0005 | | 0.00 | 0.42 |
| *Rfwd3* | Dendritic cells | 0.0005 | -2.00 | 0.36 | 0.09 |
| *Twf2* | T cells | 0.0005 | -1.39 | 1.19 | 0.45 |
| *Myh6* | Juxtaglomerular cells | 0.0005 | | 0.00 | 0.31 |

| | | | | | |
|---|---|---|---|---|---|
| *Gm39469* | Proximal tubule cells | 0.0005 | | 0.29 | 0.00 |
| *Atp5c1* | Dendritic cells | 0.0005 | -0.35 | 4.36 | 3.43 |
| *Cd2* | T cells | 0.0005 | 0.99 | 2.49 | 4.96 |
| *Phf11d* | T cells | 0.0005 | | 0.00 | 0.21 |
| *Parm1* | Collecting duct principal cells | 0.0005 | -2.13 | 3.95 | 0.90 |
| *Cox16* | Natural killer cells | 0.0005 | | 1.02 | 0.00 |
| *Sft2d3* | Collecting duct principal cells | 0.0006 | | 0.28 | 0.00 |
| *Gm21762* | Dendritic cells | 0.0006 | | 0.87 | 0.00 |
| *Fbxl7* | Collecting duct principal cells | 0.0006 | | 0.18 | 0.00 |
| *Gemin7* | Dendritic cells | 0.0007 | -1.10 | 0.63 | 0.30 |
| *Rapgef6* | B cells | 0.0007 | -0.45 | 1.78 | 1.30 |
| *C1qc* | Proximal tubule cells | 0.0007 | -0.93 | 0.59 | 0.31 |
| *Rusc1* | B cells | 0.0007 | | 0.24 | 0.00 |
| *Prpf4* | Collecting duct principal cells | 0.0007 | | 0.19 | 0.00 |
| *H3f3b* | Juxtaglomerular cells | 0.0007 | -0.75 | 27.23 | 16.21 |
| *Timm44* | T cells | 0.0007 | 1.82 | 0.26 | 0.92 |
| *Topors* | Collecting duct principal cells | 0.0007 | -1.58 | 0.85 | 0.28 |
| *Dynlt1b* | Fibroblast | 0.0007 | -3.77 | 0.60 | 0.04 |
| *Flt3l* | NK cells | 0.0008 | | 0.00 | 0.72 |
| *Cnn2* | Fibroblast | 0.0009 | -0.10 | 2.65 | 2.46 |
| *Hpf1* | NK cells | 0.0009 | 4.55 | 0.06 | 1.50 |
| *Cebpz* | Fibroblast | 0.001 | -0.16 | 1.12 | 1.00 |
| *Mrps17* | Juxtaglomerular cells | 0.001 | -1.07 | 1.38 | 0.66 |
| *Sat1* | B cells | 0.001 | 0.56 | 1.85 | 2.73 |
| *Smc6* | Juxtaglomerular cells | 0.001 | 1.91 | 0.42 | 1.56 |
| *Lrch1* | B cells | 0.001 | | 0.00 | 0.36 |
| *Siva1* | Fibroblast | 0.001 | 2.39 | 0.21 | 1.09 |
| *Rpl7a* | B cells | 0.001 | -0.48 | 14.58 | 10.46 |
| *Diaph2* | Fibroblast | 0.001 | | 0.00 | 0.66 |
| *Stk16* | Natural killer cells | 0.001 | | 0.83 | 0.00 |
| *Ilk* | Natural killer cells | 0.002 | | 0.87 | 0.00 |
| *Mapkapk3* | Fibroblast | 0.002 | -2.30 | 0.74 | 0.15 |

| | | | | | |
|---|---|---|---|---|---|
| *Vps53* | Natural killer cells | 0.002 | | 1.20 | 0.00 |
| *Nhp2* | Natural killer cells | 0.002 | -0.70 | 1.43 | 0.88 |
| *Ptrhd1* | Natural killer cells | 0.003 | | 0.00 | 0.86 |
| *Cyth1* | Natural killer cells | 0.003 | | 1.78 | 0.00 |

Legend: KO, miR-181a knockout mice; WT, wild-type mice.

**Table S3.** Up-regulated gene ontology pathways based on cell type.

| Cell type | ID | Description | GeneRatio | *P*-value | FDR q-value | Gene ID | Count |
|---|---|---|---|---|---|---|---|
| B cells | GO:0042026 | protein refolding | 4/15 | 2.8047E-08 | 1.87914E-05 | *Hspa1b/Hspd1/Hsp90aa1/Hspa1a* | 4 |
| | GO:0006986 | response to unfolded protein | 4/15 | 3.6804E-06 | 0.000616469 | *Hspa1b/Hspd1/Hsp90aa1/Hspa1a* | 4 |
| | GO:0034605 | cellular response to heat | 3/15 | 3.0901E-05 | 0.002300415 | *Hspa1b/Hsp90aa1/Hspa1a* | 3 |
| | GO:1901673 | regulation of mitotic spindle assembly | 2/15 | 0.00022228 | 0.013539153 | *Hspa1b/Hspa1a* | 2 |
| | GO:1902850 | microtubule cytoskeleton organization involved in mitosis | 3/15 | 0.00030721 | 0.016561511 | *Pafah1b1/Hspa1b/Hspa1a* | 3 |
| | GO:1903364 | positive regulation of cellular protein catabolic process | 3/15 | 0.00033805 | 0.016561511 | *Hspa1b/Hsp90aa1/Hspa1a* | 3 |
| | GO:2000778 | positive regulation of interleukin-6 secretion | 2/15 | 0.00043993 | 0.018422221 | *Hspd1/Ncl* | 2 |
| | GO:0001819 | positive regulation of cytokine production | 4/15 | 0.00060189 | 0.023721514 | *Hspd1/Ncl/Icosl/Ddx21* | 4 |
| | GO:0021955 | central nervous system neuron axonogenesis | 2/15 | 0.00068934 | 0.024308381 | *Pafah1b1/Hsp90aa1* | 2 |
| | GO:0050870 | positive regulation of T cell activation | 3/15 | 0.00073467 | 0.024611566 | *Hspd1/Icosl/Hsp90aa1* | 3 |
| | GO:1903039 | positive regulation of leukocyte cell-cell adhesion | 3/15 | 0.00093319 | 0.026072567 | *Hspd1/Icosl/Hsp90aa1* | 3 |
| | GO:0002440 | production of molecular mediator of immune response | 3/15 | 0.00101157 | 0.026072567 | *Hspd1/Icosl/Ddx21* | 3 |

| | | | | | | |
|---|---|---|---|---|---|---|
| GO:0072604 | interleukin-6 secretion | 2/15 | 0.00104075 | 0.026072567 | *Hspd1/Ncl* | 2 |
| GO:0002204 | somatic recombination of immunoglobulin genes involved in immune response | 2/15 | 0.0010896 | 0.026072567 | *Hspd1/Icosl* | 2 |
| GO:0002208 | somatic diversification of immunoglobulins involved in immune response | 2/15 | 0.0010896 | 0.026072567 | *Hspd1/Icosl* | 2 |
| GO:0045190 | isotype switching | 2/15 | 0.0010896 | 0.026072567 | *Hspd1/Icosl* | 2 |
| GO:0032886 | regulation of microtubule-based process | 3/15 | 0.00144597 | 0.031127639 | *Pafah1b1/Hspa1b/Hspa1a* | 3 |
| GO:0051251 | positive regulation of lymphocyte activation | 3/15 | 0.00203274 | 0.034048329 | *Hspd1/Icosl/Hsp90aa1* | 3 |
| GO:0006626 | protein targeting to mitochondrion | 2/15 | 0.00243552 | 0.037948828 | *Hspd1/Hsp90aa1* | 2 |
| GO:0002312 | B cell activation involved in immune response | 2/15 | 0.00258318 | 0.039334723 | *Hspd1/Icosl* | 2 |
| GO:0021954 | central nervous system neuron development | 2/15 | 0.00265856 | 0.039582965 | *Pafah1b1/Hsp90aa1* | 2 |
| GO:0034620 | cellular response to unfolded protein | 2/15 | 0.00289089 | 0.041210625 | *Hspa1b/Hspa1a* | 2 |
| GO:0007159 | leukocyte cell-cell adhesion | 3/15 | 0.00322757 | 0.044132131 | *Hspd1/Icosl/Hsp90aa1* | 3 |
| GO:0072655 | establishment of protein localization to mitochondrion | 2/15 | 0.0037318 | 0.047321181 | *Hspd1/Hsp90aa1* | 2 |
| GO:1903426 | regulation of reactive oxygen species biosynthetic process | 2/15 | 0.00391213 | 0.047321181 | *Hspd1/Hsp90aa1* | 2 |
| GO:0002443 | leukocyte mediated immunity | 3/15 | 0.00409235 | 0.047321181 | *Hspd1/Icosl/Ddx21* | 3 |

| | | | | | | | |
|---|---|---|---|---|---|---|---|
| | GO:0070585 | protein localization to mitochondrion | 2/15 | 0.00409646 | 0.047321181 | *Hspd1/Hsp90aa1* | 2 |
| Collecting duct principal cells | GO:0001702 | gastrulation with mouth forming second | 3/33 | 7.1857E-05 | 0.040334342 | *Smad4/Ldb1/Ets2* | 3 |
| | GO:0009948 | anterior/posterior axis specification | 3/33 | 8.5909E-05 | 0.040334342 | *Smad4/Ldb1/Ets2* | 3 |
| Endothelial cells | GO:0001525 | angiogenesis | 23/181 | 1.6374E-08 | 4.37684E-05 | *Rgcc/Col4a1/Sparc/Mtdh/Esm1/Rhoa/Flt1/Mmrn2/Hspb1/Col4a2/Pecam1/Mfge8/Ptprb/Aplnr/Myh9/Wasf2/Ptprm/Tcf4/Ets1/Casp8/B4galt1/Sema6a/Tgfbr2* | 23 |
| | GO:0035455 | response to interferon-alpha | 5/181 | 7.477E-06 | 0.006662036 | *Ifit3/Tpr/Bst2/Ifitm2/Ifit1* | 5 |
| | GO:0001667 | ameboidal-type cell migration | 16/181 | 2.2218E-05 | 0.008825877 | *Rgcc/Sparc/Calr/Rhoa/Mmrn2/Hspb1/Pecam1/Scarb1/Myh9/Wasf2/Ptprm/Ets1/Sema3d/Sema3f/Sema6a/Tgfbr2* | 16 |
| | GO:0043542 | endothelial cell migration | 11/181 | 2.6776E-05 | 0.008825877 | *Rgcc/Sparc/Calr/Rhoa/Mmrn2/Hspb1/Pecam1/Scarb1/Myh9/Ptprm/Ets1* | 11 |
| | GO:0002486 | antigen processing and presentation of endogenous peptide antigen via MHC class I via ER pathway, TAP-independent | 4/181 | 2.9682E-05 | 0.008825877 | *H2-D1/H2-K1/H2-Q6/H2-Q7* | 4 |
| | GO:0043062 | extracellular structure organization | 13/181 | 3.4137E-05 | 0.008825877 | *Rgcc/Col4a1/App/Col15a1/Col4a2/Vwa1/Scarb1/Serpinh1/Ets1/Abcg1/Fscn1/B4galt1/Gpihbp1* | 13 |

| | | | | | | |
|---|---|---|---|---|---|---|
| GO:0002476 | antigen processing and presentation of endogenous peptide antigen via MHC class Ib | 4/181 | 4.0046E-05 | 0.008825877 | *H2-D1/H2-K1/H2-Q6/H2-Q7* | 4 |
| GO:0002484 | antigen processing and presentation of endogenous peptide antigen via MHC class I via ER pathway | 4/181 | 4.0046E-05 | 0.008825877 | *H2-D1/H2-K1/H2-Q6/H2-Q7* | 4 |
| GO:0002474 | antigen processing and presentation of peptide antigen via MHC class I | 5/181 | 4.7419E-05 | 0.008825877 | *H2-D1/H2-K1/H2-Q6/Calr/H2-Q7* | 5 |
| GO:0002428 | antigen processing and presentation of peptide antigen via MHC class Ib | 4/181 | 5.283E-05 | 0.008825877 | *H2-D1/H2-K1/H2-Q6/H2-Q7* | 4 |
| GO:1900151 | regulation of nuclear-transcribed mRNA catabolic process, deadenylation-dependent decay | 4/181 | 5.283E-05 | 0.008825877 | *Zfp36l1/Btg2/Tnrc6c/Tnrc6b* | 4 |
| GO:1900153 | positive regulation of nuclear-transcribed mRNA catabolic process, deadenylation-dependent decay | 4/181 | 5.283E-05 | 0.008825877 | *Zfp36l1/Btg2/Tnrc6c/Tnrc6b* | 4 |
| GO:0001937 | negative regulation of endothelial cell proliferation | 5/181 | 8.8605E-05 | 0.01393188 | *Rgcc/Sparc/Flt1/Ptprm/Gja1* | 5 |
| GO:0010594 | regulation of endothelial cell migration | 9/181 | 9.8619E-05 | 0.014528164 | *Rgcc/Sparc/Calr/Rhoa/Mmrn2/Hspb1/Ptprm/Ets1/Plpp3* | 9 |
| GO:0090130 | tissue migration | 12/181 | 0.00014696 | 0.016936007 | *Rgcc/Sparc/Calr/Rhoa/Mmrn2/Hspb1/Pecam1/Scarb1/Myh9/Ptprm/Ets1/Tgfbr2* | 12 |
| GO:0090132 | epithelium migration | 12/181 | 0.00014696 | 0.016936007 | *Rgcc/Sparc/Calr/Rhoa/Mmrn2/Hspb1/Pecam1/Scarb1/Myh9/Ptprm/Ets1/Tgfbr2* | 12 |

| | | | | | | |
|---|---|---|---|---|---|---|
| GO:0060348 | bone development | 10/181 | 0.00015912 | 0.016936007 | *Sparc/Rflnb/Rhoa/Vwa1/Serpinh1/Hoxb4/Wasf2/Fli1/Gja1/Tgfbr2* | 10 |
| GO:0061572 | actin filament bundle organization | 9/181 | 0.00025305 | 0.024156921 | *Rgcc/Hsp90b1/Rflnb/Rdx/Limch1/Rhoa/Wasf2/Cald1/Fscn1* | 9 |
| GO:0070848 | response to growth factor | 17/181 | 0.00029393 | 0.027092266 | *Sparc/Ltbp4/Flt1/Mmrn2/Zfhx3/Hspb1/Col4a2/Ltbp1/Hsp90ab1/Tcf4/Rab14/Tpr/Ier2/Zfp36l1/Sema6a/Ehd4/Tgfbr2* | 17 |
| GO:0035457 | cellular response to interferon-alpha | 3/181 | 0.00037579 | 0.030438902 | *Ifit3/Tpr/Ifit1* | 3 |
| GO:0031331 | positive regulation of cellular catabolic process | 13/181 | 0.00043247 | 0.033053081 | *Mtdh/Rdx/Igfbp3/Wac/Msn/Qk/Rnf144a/Zfp36l1/Pttg1ip/Btg2/Tnrc6c/Tnrc6b/Trib2* | 13 |
| GO:0040013 | negative regulation of locomotion | 12/181 | 0.00043279 | 0.033053081 | *Rgcc/Calr/Igfbp3/Limch1/Cd200/Rhoa/Mmrn2/Ptprm/Trim56/Sema3d/Sema3f/Sema6a* | 12 |
| GO:0034260 | negative regulation of GTPase activity | 4/181 | 0.00045327 | 0.033655314 | *Rdx/Pecam1/Rasa4/Wnk1* | 4 |
| GO:0071803 | positive regulation of podosome assembly | 3/181 | 0.00049605 | 0.033993715 | *Rhoa/Msn/Fscn1* | 3 |
| GO:0001916 | positive regulation of T cell mediated cytotoxicity | 4/181 | 0.00052324 | 0.033993715 | *H2-D1/H2-K1/H2-Q6/H2-Q7* | 4 |
| GO:0006457 | protein folding | 8/181 | 0.00061754 | 0.037569699 | *Hsp90b1/Pdia3/Calr/Hspb1/Hsp90ab1/Canx/Unc45b/Ppic* | 8 |

| | | | | | | |
|---|---|---|---|---|---|---|
| GO:0009896 | positive regulation of catabolic process | 14/181 | 0.00067401 | 0.038929532 | *Mtdh/Rdx/Igfbp3/Wac/Msn/Qk/Rnf144a/Zfp36l1/Pttg1ip/Btg2/Tnrc6c/Gja1/Tnrc6b/Trib2* | 14 |
| GO:0006417 | regulation of translation | 12/181 | 0.000689 | 0.038929532 | *Calr/App/Gtpbp2/Qk/Tpr/Trnau1ap/Zfp36l1/Btg2/Tnrc6c/Tnrc6b/Nck1/Magoh* | 12 |
| GO:0035304 | regulation of protein dephosphorylation | 6/181 | 0.00070855 | 0.038929532 | *Hsp90b1/Hsp90ab1/Ppp2r5a/Ikbkb/Ywhab/Ppp1r12a* | 6 |
| GO:0010563 | negative regulation of phosphorus metabolic process | 16/181 | 0.00079844 | 0.040035959 | *Igfbp3/Rhoa/Hspb1/Pecam1/Sh3bp5/Entpd1/Fabp4/Plpp3/Gnaq/Ppp2r5a/Ikbkb/Dynll1/Ywhab/Sema6a/Nck1/Wnk1* | 16 |
| GO:0045936 | negative regulation of phosphate metabolic process | 16/181 | 0.00079844 | 0.040035959 | *Igfbp3/Rhoa/Hspb1/Pecam1/Sh3bp5/Entpd1/Fabp4/Plpp3/Gnaq/Ppp2r5a/Ikbkb/Dynll1/Ywhab/Sema6a/Nck1/Wnk1* | 16 |
| GO:0034368 | protein-lipid complex remodeling | 3/181 | 0.00080447 | 0.040035959 | *Scarb1/Abcg1/Gpihbp1* | 3 |
| GO:0034369 | plasma lipoprotein particle remodeling | 3/181 | 0.00080447 | 0.040035959 | *Scarb1/Abcg1/Gpihbp1* | 3 |
| GO:1901264 | carbohydrate derivative transport | 5/181 | 0.0009558 | 0.04404911 | *Slc29a1/Scarb1/Abcg1/Gja1/Ralbp1* | 5 |
| GO:0035089 | establishment of apical/basal cell polarity | 3/181 | 0.00121313 | 0.049887561 | *Rhoa/Msn/Fscn1* | 3 |

| | | | | | | | |
|---|---|---|---|---|---|---|---|
| Macrophages | GO:0019886 | antigen processing and presentation of exogenous peptide antigen via MHC class II | 4/82 | 9.1443E-07 | 0.00199163 | *H2-Aa/H2-DMa/H2-DMb1/H2-Ab1* | 4 |
| | GO:0055094 | response to lipoprotein particle | 4/82 | 0.000015293 | 0.00555151 | *Cd81/Apoe/Cd36/Itgb2* | 4 |
| | GO:0071402 | cellular response to lipoprotein particle stimulus | 4/82 | 0.00002102 | 0.00654008 | *Cd81/Apoe/Cd36/Itgb2* | 4 |
| | GO:0016042 | lipid catabolic process | 8/82 | 0.000051732 | 0.01251913 | *Apoe/Pla2g7/Psap/Plbd1/Lipe/Smpdl3a/Pld4/Daglb* | 8 |
| | GO:2000377 | regulation of reactive oxygen species metabolic process | 7/82 | 0.000092537 | 0.01920624 | *Cd36/Cx3cr1/Itgb2/Prcp/Ogt/Pid1/Tspo* | 7 |
| | GO:0042136 | neurotransmitter biosynthetic process | 5/82 | 0.00013885 | 0.02244849 | *Cd36/Cx3cr1/Itgb2/Tspo/Daglb* | 5 |
| | GO:0046339 | diacylglycerol metabolic process | 3/82 | 0.00014369 | 0.02244849 | *Lipe/Ang/Daglb* | 3 |
| | GO:0006898 | receptor-mediated endocytosis | 7/82 | 0.00015536 | 0.02244849 | *Trf/Cd81/Apoe/Cd36/Ap2a2/Itgb2/Fcho2* | 7 |
| | GO:0071222 | cellular response to lipopolysaccharide | 6/82 | 0.00027626 | 0.02865253 | *Cd36/Cx3cr1/Cd84/Sbno2/Ogt/Ly86* | 6 |
| | GO:1903409 | reactive oxygen species biosynthetic process | 5/82 | 0.00029359 | 0.02888192 | *Cd36/Cx3cr1/Itgb2/Ogt/Tspo* | 5 |
| | GO:0006959 | humoral immune response | 5/82 | 0.00040812 | 0.0341883 | *Cd81/C1qc/H2-DMa/Cfh/H2-Ab1* | 5 |
| | GO:0032387 | negative regulation of intracellular transport | 4/82 | 0.00043963 | 0.03496405 | *Cd36/Snx3/Cd84/Ogt* | 4 |

| | | | | | | |
|---|---|---|---|---|---|---|
| GO:0002443 | leukocyte mediated immunity | 8/82 | 0.00047422 | 0.03496405 | *Cd81/H2-D1/Ctsh/Itgb2/C1qc/Cd84/H2-DMa/H2-Ab1* | 8 |
| GO:0071396 | cellular response to lipid | 9/82 | 0.00048268 | 0.03496405 | *Cd36/Cx3cr1/Pdia3/Cd84/Sbno2/Ogt/Pid1/Ep300/Ly86* | 9 |
| GO:0072657 | protein localization to membrane | 10/82 | 0.00051371 | 0.03496405 | *Cd81/Apoe/Zmynd8/Itgb2/Timm13/Fcho2/Tnfrsf1a/Ogt/Pid1/Myadm* | 10 |
| GO:0002521 | leukocyte differentiation | 10/82 | 0.0005495 | 0.03626717 | *Trf/Cd81/Csf1r/Psen2/H2-Aa/C1qc/Sbno2/H2-DMa/Ep300/H2-Ab1* | 10 |
| GO:0097242 | amyloid-beta clearance | 3/82 | 0.00065924 | 0.04102358 | *Apoe/Cd36/Itgb2* | 3 |
| GO:0071216 | cellular response to biotic stimulus | 6/82 | 0.00070248 | 0.04212341 | *Cd36/Cx3cr1/Cd84/Sbno2/Ogt/Ly86* | 6 |
| GO:0006809 | nitric oxide biosynthetic process | 4/82 | 0.00071949 | 0.04212341 | *Cd36/Cx3cr1/Itgb2/Tspo* | 4 |
| GO:0051092 | positive regulation of NF-kappaB transcription factor activity | 5/82 | 0.00073494 | 0.04212341 | *Rab7b/Cd36/Cx3cr1/Itgb2/Cd84* | 5 |
| GO:0045765 | regulation of angiogenesis | 7/82 | 0.00080703 | 0.04435691 | *Cd36/Ctsh/Cx3cr1/Itgb2/Glul/Tnfrsf1a/Krit1* | 7 |
| GO:0050778 | positive regulation of immune response | 10/82 | 0.00093179 | 0.04800003 | *Cd81/H2-D1/Psen2/Rab7b/Cd36/Itgb2/C1qc/H2-DMa/Cfh/H2-Ab1* | 10 |

| | | | | | | | |
|---|---|---|---|---|---|---|---|
| | GO:0010324 | membrane invagination | 4/82 | 0.00094766 | 0.04800003 | *Cd36/Itgb2/Snx3/Fcho2* | 4 |
| | GO:0002275 | myeloid cell activation involved in immune response | 4/82 | 0.00099875 | 0.04833933 | *Cx3cr1/Itgb2/Cd84/Sbno2* | 4 |
| | GO:2001057 | reactive nitrogen species metabolic process | 4/82 | 0.00099875 | 0.04833933 | *Cd36/Cx3cr1/Itgb2/Tspo* | 4 |
| T cells | GO:0002521 | leukocyte differentiation | 7/27 | 0.00002922 2 | 0.01349095 | *Ccl5/Cd8a/Ptprc/Lgals1/Runx3/Tn faip3/Tox* | 7 |
| | GO:0043900 | regulation of multi-organism process | 6/27 | 0.00003735 7 | 0.01349095 | *Ccl5/Hspa8/Lgals1/Tpst2/Tnfaip3/ Ccl4* | 6 |
| | GO:0030098 | lymphocyte differentiation | 6/27 | 0.00003934 3 | 0.01349095 | *Cd8a/Ptprc/Lgals1/Runx3/Tnfaip3 /Tox* | 6 |
| | GO:0051249 | regulation of lymphocyte activation | 6/27 | 0.00007496 6 | 0.01349095 | *Ccl5/Ptprc/Lgals1/Runx3/Tnfaip3/ Tox* | 6 |
| | GO:2001233 | regulation of apoptotic signaling pathway | 6/27 | 0.00007726 4 | 0.01349095 | *Birc6/Ptprc/Triap1/Dnaja1/Runx3 /Tnfaip3* | 6 |
| | GO:0051251 | positive regulation of lymphocyte activation | 5/27 | 0.00009062 3 | 0.01349095 | *Ccl5/Ptprc/Lgals1/Runx3/Tox* | 5 |
| | GO:0034113 | heterotypic cell-cell adhesion | 3/27 | 0.00009307 1 | 0.01349095 | *Cd2/Ptprc/Tnfaip3* | 3 |
| | GO:0042110 | T cell activation | 6/27 | 0.00013823 | 0.01349095 | *Ccl5/Cd2/Cd8a/Ptprc/Lgals1/Run x3* | 6 |

| | | | | | | |
|---|---|---|---|---|---|---|
| GO:2001236 | regulation of extrinsic apoptotic signaling pathway | 4/27 | 0.00016539 | 0.01349095 | *Birc6/Ptprc/Runx3/Tnfaip3* | 4 |
| GO:0043901 | negative regulation of multi-organism process | 4/27 | 0.00017002 | 0.01349095 | *Ccl5/Tpst2/Tnfaip3/Ccl4* | 4 |
| GO:0032680 | regulation of tumor necrosis factor production | 4/27 | 0.00017475 | 0.01349095 | *Cd2/Ptprc/Tnfaip3/Ccl4* | 4 |
| GO:1903555 | regulation of tumor necrosis factor superfamily cytokine production | 4/27 | 0.00018447 | 0.01349095 | *Cd2/Ptprc/Tnfaip3/Ccl4* | 4 |
| GO:0032640 | tumor necrosis factor production | 4/27 | 0.00018948 | 0.01349095 | *Cd2/Ptprc/Tnfaip3/Ccl4* | 4 |
| GO:1902107 | positive regulation of leukocyte differentiation | 4/27 | 0.00018948 | 0.01349095 | *Ccl5/Ptprc/Runx3/Tox* | 4 |
| GO:0071706 | tumor necrosis factor superfamily cytokine production | 4/27 | 0.00019979 | 0.01349095 | *Cd2/Ptprc/Tnfaip3/Ccl4* | 4 |
| GO:0051851 | modification by host of symbiont morphology or physiology | 3/27 | 0.00020204 | 0.01349095 | *Ccl5/Hspa8/Ccl4* | 3 |
| GO:0002696 | positive regulation of leukocyte activation | 5/27 | 0.00020459 | 0.01349095 | *Ccl5/Ptprc/Lgals1/Runx3/Tox* | 5 |
| GO:0051702 | interaction with symbiont | 3/27 | 0.00024717 | 0.01352395 | *Ccl5/Hspa8/Ccl4* | 3 |
| GO:0048245 | eosinophil chemotaxis | 2/27 | 0.00030442 | 0.01484135 | *Ccl5/Ccl4* | 2 |
| GO:0050870 | positive regulation of T cell activation | 4/27 | 0.00030451 | 0.01484135 | *Ccl5/Ptprc/Lgals1/Runx3* | 4 |
| GO:0002312 | B cell activation involved in immune response | 3/27 | 0.00034095 | 0.01528835 | *Ptprc/Lgals1/Tnfaip3* | 3 |
| GO:1903039 | positive regulation of leukocyte cell-cell adhesion | 4/27 | 0.00041649 | 0.01703441 | *Ccl5/Ptprc/Lgals1/Runx3* | 4 |

| | | | | | | |
|---|---|---|---|---|---|---|
| GO:0072677 | eosinophil migration | 2/27 | 0.00046659 | 0.01743485 | *Ccl5/Ccl4* | 2 |
| GO:0140131 | positive regulation of lymphocyte chemotaxis | 2/27 | 0.00046659 | 0.01743485 | *Ccl5/Ccl4* | 2 |
| GO:0032897 | negative regulation of viral transcription | 2/27 | 0.00073508 | 0.02112882 | *Ccl5/Ccl4* | 2 |
| GO:1901623 | regulation of lymphocyte chemotaxis | 2/27 | 0.00073508 | 0.02112882 | *Ccl5/Ccl4* | 2 |
| GO:0002456 | T cell mediated immunity | 3/27 | 0.00099371 | 0.02370099 | *Cd8a/Hspa8/Ptprc* | 3 |
| GO:0043270 | positive regulation of ion transport | 4/27 | 0.00109446 | 0.02556026 | *Ccl5/Triap1/Slc25a4/Ccl4* | 4 |
| GO:0016032 | viral process | 4/27 | 0.00144767 | 0.02798394 | *Ccl5/Hspa8/Lgals1/Ccl4* | 4 |
| GO:0001916 | positive regulation of T cell mediated cytotoxicity | 2/27 | 0.00144788 | 0.02798394 | *Hspa8/Ptprc* | 2 |
| GO:0035025 | positive regulation of Rho protein signal transduction | 2/27 | 0.00144788 | 0.02798394 | *P2ry10/Akap13* | 2 |
| GO:0010667 | negative regulation of cardiac muscle cell apoptotic process | 2/27 | 0.00155318 | 0.02901867 | *Hspa8/Slc25a4* | 2 |
| GO:0072666 | establishment of protein localization to vacuole | 2/27 | 0.00238914 | 0.03997359 | *Hspa8/Tnfaip3* | 2 |
| GO:0003298 | physiological muscle hypertrophy | 2/27 | 0.00252252 | 0.04039631 | *Akap13/Slc25a4* | 2 |
| GO:0003301 | physiological cardiac muscle hypertrophy | 2/27 | 0.00252252 | 0.04039631 | *Akap13/Slc25a4* | 2 |
| GO:0061049 | cell growth involved in cardiac muscle cell development | 2/27 | 0.00252252 | 0.04039631 | *Akap13/Slc25a4* | 2 |

| | | | | | | | |
|---|---|---|---|---|---|---|---|
| | GO:0019932 | second-messenger-mediated signaling | 4/27 | 0.00256917 | 0.04056388 | *Cd8a/Ptprc/Akap13/Ccl4* | 4 |
| | GO:0070098 | chemokine-mediated signaling pathway | 2/27 | 0.00279959 | 0.04240999 | *Ccl5/Ccl4* | 2 |

**Table S4.** Down-regulated gene ontology pathways based on cell type.

| Cell type | ID | Description | GeneRatio | *P*-value | FDR q-value | Gene ID | Count |
|---|---|---|---|---|---|---|---|
| Dendritic cells | GO:0000226 | microtubule cytoskeleton organization | 9/43 | 1.7678E-05 | 0.01225581 | *Dynlt1b/Cep295/Ccnb1/Bora/Sgo1/Dync1li2/Tuba1c/Cnp/Birc5* | 9 |
| | GO:0090068 | positive regulation of cell cycle process | 6/43 | 5.3577E-05 | 0.01225581 | *Cep295/Ccnb1/E2f8/Ddx3x/Ube2c/Birc5* | 6 |
| | GO:0051303 | establishment of chromosome localization | 4/43 | 5.5243E-05 | 0.01225581 | *Ccnb1/Cenpf/Cdca8/Birc5* | 4 |
| | GO:0050000 | chromosome localization | 4/43 | 5.867E-05 | 0.01225581 | *Ccnb1/Cenpf/Cdca8/Birc5* | 4 |
| | GO:0044772 | mitotic cell cycle phase transition | 7/43 | 7.1663E-05 | 0.01225581 | *Rfwd3/Ccnb1/Ccna2/Cenpf/Ddx3x/Ube2c/Birc5* | 7 |
| | GO:1901990 | regulation of mitotic cell cycle phase transition | 6/43 | 9.0864E-05 | 0.01225581 | *Rfwd3/Ccnb1/Cenpf/Ddx3x/Ube2c/Birc5* | 6 |
| | GO:1901992 | positive regulation of mitotic cell cycle phase transition | 4/43 | 9.6881E-05 | 0.01225581 | *Ccnb1/Ddx3x/Ube2c/Birc5* | 4 |
| | GO:0140014 | mitotic nuclear division | 6/43 | 9.7559E-05 | 0.01225581 | *Ccnb1/Cdca8/Bora/Sgo1/Ube2c/Birc5* | 6 |

| | | | | | | | |
|---|---|---|---|---|---|---|---|
| | GO:0050685 | positive regulation of mRNA processing | 3/43 | 0.00015981 | 0.0138158 | *Ccnb1/Sf3b4/Dazap1* | 3 |
| | GO:0000280 | nuclear division | 6/43 | 0.00058298 | 0.0305583 | *Ccnb1/Cdca8/Bora/Sgo1/Ube2c/Birc5* | 6 |
| | GO:0007096 | regulation of exit from mitosis | 2/43 | 0.00065822 | 0.03150056 | *Ube2c/Birc5* | 2 |
| | GO:0000070 | mitotic sister chromatid segregation | 4/43 | 0.00099503 | 0.041667 | *Ccnb1/Cdca8/Sgo1/Birc5* | 4 |
| | GO:0008625 | extrinsic apoptotic signaling pathway via death domain receptors | 3/43 | 0.00130007 | 0.04652476 | *Gpx1/Ddx3x/Psen2* | 3 |
| | GO:0006379 | mRNA cleavage | 2/43 | 0.00134251 | 0.04652476 | *Polr2i/Cstf2* | 2 |
| | GO:0043154 | negative regulation of cysteine-type endopeptidase activity involved in apoptotic process | 3/43 | 0.00141553 | 0.04742025 | *Gpx1/Ddx3x/Birc5* | 3 |
| Endothelial cells | GO:0002181 | cytoplasmic translation | 18/175 | 1.2508E-17 | 2.6666E-14 | *Rpl13a/Rps28/Rpl6/Rps26/Rpl17/Rpl39/Rpl35a/Rpl36/Rpl18a/Rpl10a/Rpl9/Rpl24/Rpl30/Mcts1/Rps23/Eef2/Rpsa/Rpl18* | 18 |
| | GO:0022613 | ribonucleoprotein complex biogenesis | 24/175 | 3.7003E-10 | 3.9446E-07 | *Rpl13a/Rps28/Rpl6/Rps19/Rpl35/Pin4/Rps27/Rpl14/Rpl27/Nop10/Rpl35a/Rpl10a/Snrpb/Rpl24/Snrpd1/Rps* | 24 |

| | | | | | | |
|---|---|---|---|---|---|---|
| | | | | | *10/Rpl7/Sf3a2/Mcts1/Rps16/Rps23/Rps5/Rpsa/Rpl12* | |
| GO:0033108 | mitochondrial respiratory chain complex assembly | 11/175 | 3.4737E-09 | 1.9326E-06 | *Ndufb9/Ndufs5/Ndufa5/Uqcrb/Ndufs8/Ndufs7/Ndufa2/Ndufa6/Ndufa3/Ndufa1/Cox17* | 11 |
| GO:0010257 | NADH dehydrogenase complex assembly | 9/175 | 4.5324E-09 | 1.9326E-06 | *Ndufb9/Ndufs5/Ndufa5/Ndufs8/Ndufs7/Ndufa2/Ndufa6/Ndufa3/Ndufa1* | 9 |
| GO:0032981 | mitochondrial respiratory chain complex I assembly | 9/175 | 4.5324E-09 | 1.9326E-06 | *Ndufb9/Ndufs5/Ndufa5/Ndufs8/Ndufs7/Ndufa2/Ndufa6/Ndufa3/Ndufa1* | 9 |
| GO:0009141 | nucleoside triphosphate metabolic process | 17/175 | 1.8273E-08 | 5.5654E-06 | *Atp5l/Cox7a2/Nme2/Cox4i1/Aldoa/Ndufb9/Ndufs6/Gapdh/Atp5h/Cox7c/Uqcrb/Sdhc/Ndufs8/Suclg1/Cox6a1/Atp5g2/Atp5d* | 17 |
| GO:0009060 | aerobic respiration | 9/175 | 5.815E-08 | 1.0331E-05 | *Cox4i1/Cox7c/Uqcrb/Sdhc/Ndufs8/Suclg1/Ndufs7/Cox6a1/Atp5d* | 9 |
| GO:0022618 | ribonucleoprotein complex assembly | 15/175 | 7.1038E-08 | 1.165E-05 | *Rpl13a/Rps28/Rpl6/Rps19/Rps27/Snrpb/Rpl24/Snrpd1/Rps10/Sf3a2/Mcts1/Rps23/Rps5/Rpsa/Rpl12* | 15 |
| GO:0071826 | ribonucleoprotein complex subunit organization | 15/175 | 1.4223E-07 | 1.8952E-05 | *Rpl13a/Rps28/Rpl6/Rps19/Rps27/Snrpb/Rpl24/Snrpd1/Rps10/Sf3a2/Mcts1/Rps23/Rps5/Rpsa/Rpl12* | 15 |

| | | | | | | |
|---|---|---|---|---|---|---|
| GO:0019693 | ribose phosphate metabolic process | 18/175 | 1.941E-06 | 0.00013794 | *Atp5l/Cox7a2/Nme2/Cox4i1/Aldoa/Ndufb9/Ndufs6/Gapdh/Hint1/Atp5h/Cox7c/Uqcrb/Sdhc/Ndufs8/Suclg1/Cox6a1/Atp5g2/Atp5d* | 18 |
| GO:0006091 | generation of precursor metabolites and energy | 16/175 | 2.136E-06 | 0.0001469 | *Cox7a2/Cox4i1/Aldoa/Ndufb9/Ndufs6/Gapdh/Cox7c/Ndufa5/Uqcrb/Sdhc/Ndufs8/Suclg1/Ndufs7/Cox6a1/Atp5d/Cox17* | 16 |
| GO:0072521 | purine-containing compound metabolic process | 18/175 | 5.186E-06 | 0.00033505 | *Atp5l/Cox7a2/Nme2/Cox4i1/Aldoa/Ndufb9/Ndufs6/Gapdh/Hint1/Atp5h/Cox7c/Uqcrb/Sdhc/Ndufs8/Suclg1/Cox6a1/Atp5g2/Atp5d* | 18 |
| GO:0061844 | antimicrobial humoral immune response mediated by antimicrobial peptide | 5/175 | 1.2833E-05 | 0.0008047 | *Rps19/Rpl39/Gapdh/Fau/Rpl30* | 5 |
| GO:0006364 | rRNA processing | 11/175 | 2.0634E-05 | 0.00122198 | *Rps28/Rps19/Rpl35/Pin4/Rpl14/Rpl27/Nop10/Rpl35a/Rpl10a/Rpl7/Rps16* | 11 |
| GO:0007005 | mitochondrion organization | 17/175 | 4.7461E-05 | 0.00256176 | *Cox7a2/Gabarap/Ndufb9/Ndufs6/Ndufs5/Ndufa5/Uqcrb/Ndufs8/Ndufs7/Ndufa2/Ndufa6/Slc25a5/Ndufa3/Atp5d/Grpel1/Ndufa1/Cox17* | 17 |

| | | | | | | | |
|---|---|---|---|---|---|---|---|
| | GO:0015985 | energy coupled proton transport, down electrochemical gradient | 4/175 | 5.9944E-05 | 0.00304288 | *Atp5l/Atp5h/Atp5g2/Atp5d* | 4 |
| | GO:0015986 | ATP synthesis coupled proton transport | 4/175 | 5.9944E-05 | 0.00304288 | *Atp5l/Atp5h/Atp5g2/Atp5d* | 4 |
| | GO:0019730 | antimicrobial humoral response | 5/175 | 0.00011462 | 0.00555401 | *Rps19/Rpl39/Gapdh/Fau/Rpl30* | 5 |
| | GO:0010499 | proteasomal ubiquitin-independent protein catabolic process | 4/175 | 0.00020971 | 0.00971938 | *Psma7/Psmb10/Psmb3/Psmb8* | 4 |
| Juxtaglomerular cells | GO:0032733 | positive regulation of interleukin-10 production | 3/23 | 1.5959E-05 | 0.01157013 | *Pibf1/Cd34/Hspd1* | 3 |
| | GO:0032613 | interleukin-10 production | 3/23 | 6.9397E-05 | 0.01677103 | *Pibf1/Cd34/Hspd1* | 3 |
| Macrophages | GO:0002181 | cytoplasmic translation | 29/148 | 5.99E-37 | 9.9793E-34 | *Rpl13a/Rpl9/Rps28/Rpl35a/Rps29/Rpl30/Rps23/Rpl8/Rpl39/Rpl18/Rpl19/Rpl17/Rpl36/Rplp1/Rpl26/Rpl38/Rpl18a/Rpl6/Rps26/Rpl24/Eef2/Rpsa/Rpl10a/Rpl29/Rpl11/Rps21/Rpl9-ps6/Gm10073/Eif4b* | 29 |
| | GO:0042254 | ribosome biogenesis | 32/148 | 1.1372E-24 | 9.4729E-22 | *Rps27/Rps19/Rps6/Rps28/Rpl35a/Pin4/Rps10/Rpl23a/Rpl5/Rps16/Rpl7/Rpl14/Rpl26/Rpl38/Rpl27/Rps14/Rpl6/Rpl3/Rpl24/Rpl12/Npm1/Rpsa/* | 32 |

| | | | | | | |
|---|---|---|---|---|---|---|
| | | | | | *Rpl10a/Rpl11/Rps21/Rpl7a/Rps24/Rps7/Rpl10/Rps17/Rpl35/Rps15* | |
| GO:0042255 | ribosome assembly | 16/148 | 3.1224E-18 | 1.0404E-15 | *Rps27/Rps19/Rps28/Rps10/Rpl23a/Rpl5/Rpl38/Rps14/Rpl6/Rpl3/Rpl24/Rpl12/Rpsa/Rpl11/Rpl10/Rps15* | 16 |
| GO:0006364 | rRNA processing | 22/148 | 2.7875E-17 | 7.74E-15 | *Rps19/Rps6/Rps28/Rpl35a/Pin4/Rpl5/Rps16/Rpl7/Rpl14/Rpl26/Rpl27/Rps14/Npm1/Rpl10a/Rpl11/Rps21/Rpl7a/Rps24/Rps7/Rps17/Rpl35/Rps15* | 22 |
| GO:0071826 | ribonucleoprotein complex subunit organization | 19/148 | 2.5042E-12 | 3.7927E-10 | *Rpl13a/Rps27/Rps19/Rps28/Rps23/Rps10/Rpl23a/Rpl5/Rpl38/Rps14/Rpl6/Rpl3/Rpl24/Rpl12/Rpsa/Rpl11/Eif4b/Rpl10/Rps15* | 19 |
| GO:0006417 | regulation of translation | 16/148 | 1.6796E-07 | 1.8654E-05 | *Rpl13a/Rps3/Rps3a1/Rps9/Rpl5/Rplp1/Rpl26/Rpl38/Eef2/Npm1/Rpl22/Rbm3/Zfp36l2/Eif3k/Rpl10/Gapdh* | 16 |
| GO:0015985 | energy coupled proton transport, down electrochemical gradient | 5/148 | 8.7946E-07 | 8.1399E-05 | *Atp5l/Atp5e/Atp5a1/Atp5h/Atp5c1* | 5 |
| GO:0015986 | ATP synthesis coupled proton transport | 5/148 | 8.7946E-07 | 8.1399E-05 | *Atp5l/Atp5e/Atp5a1/Atp5h/Atp5c1* | 5 |

| | | | | | | |
|---|---|---|---|---|---|---|
| GO:0034248 | regulation of cellular amide metabolic process | 16/148 | 9.6857E-07 | 8.4928E-05 | *Rpl13a/Rps3/Rps3a1/Rps9/Rpl5/Rplp1/Rpl26/Rpl38/Eef2/Npm1/Rpl22/Rbm3/Zfp36l2/Eif3k/Rpl10/Gapdh* | 16 |
| GO:1901798 | positive regulation of signal transduction by p53 class mediator | 5/148 | 3.5786E-06 | 0.0002981 | *Rpl23/Rpl26/Rpl11/Ubb/Rps7* | 5 |
| GO:0061844 | antimicrobial humoral immune response mediated by antimicrobial peptide | 5/148 | 5.6734E-06 | 0.00041095 | *Rps19/Fau/Rpl30/Rpl39/Gapdh* | 5 |
| GO:0010608 | posttranscriptional regulation of gene expression | 16/148 | 7.1212E-06 | 0.00049433 | *Rpl13a/Rps3/Rps3a1/Rps9/Rpl5/Rplp1/Rpl26/Rpl38/Eef2/Npm1/Rpl22/Rbm3/Zfp36l2/Eif3k/Rpl10/Gapdh* | 16 |
| GO:0051444 | negative regulation of ubiquitin-protein transferase activity | 4/148 | 2.4007E-05 | 0.00159985 | *Rpl23/Rpl5/Rpl11/Rps7* | 4 |
| GO:0019730 | antimicrobial humoral response | 5/148 | 5.1796E-05 | 0.00319603 | *Rps19/Fau/Rpl30/Rpl39/Gapdh* | 5 |
| GO:2001244 | positive regulation of intrinsic apoptotic signaling pathway | 6/148 | 6.2607E-05 | 0.00365413 | *Rps3/Rpl26/Rpl11/Ubb/Rps7/Serinc3* | 6 |
| GO:0009205 | purine ribonucleoside triphosphate metabolic process | 10/148 | 0.00017887 | 0.00903018 | *Nme2/Atp5l/Atp5e/Cox4i1/Uqcrh/Atp5a1/Atp5h/Atp6v1a/Gapdh/Atp5c1* | 10 |
| GO:0034101 | erythrocyte homeostasis | 7/148 | 0.00037874 | 0.01371698 | *Rps19/Rps6/Rps14/B2m/Klf2/Rps24/Rps17* | 7 |

| | | | | | | |
|---|---|---|---|---|---|---|
| GO:0045116 | protein neddylation | 3/148 | 0.00055451 | 0.01847636 | *Rpl5/Rpl11/Ube2f* | 3 |
| GO:0019693 | ribose phosphate metabolic process | 12/148 | 0.00090849 | 0.02655343 | *Nme2/Atp5l/Atp5e/Cox4i1/Uqcrh/Atp5a1/Hint1/Atp5h/Tkt/Atp6v1a/Gapdh/Atp5c1* | 12 |
| GO:0006959 | humoral immune response | 6/148 | 0.00095353 | 0.02738921 | *Rps19/Fau/Rpl30/Rpl39/Ptpn6/Gapdh* | 6 |
| GO:0072521 | purine-containing compound metabolic process | 12/148 | 0.00165158 | 0.04023193 | *Nme2/Atp5l/Atp5e/Cox4i1/Uqcrh/Atp5a1/Hint1/Oas1a/Atp5h/Atp6v1a/Gapdh/Atp5c1* | 12 |
| GO:2000059 | negative regulation of ubiquitin-dependent protein catabolic process | 4/148 | 0.00166222 | 0.04023193 | *Rpl23/Rpl5/Rpl11/Rps7* | 4 |
| GO:0032069 | regulation of nuclease activity | 3/148 | 0.00177288 | 0.04219456 | *Rps3/Npm1/Oas1a* | 3 |
| GO:0050821 | protein stabilization | 7/148 | 0.00184617 | 0.04332008 | *Rpl23/Rpl5/Npm1/Rpl11/Rps7/Cct4/Gapdh* | 7 |
| GO:0031640 | killing of cells of other organism | 3/148 | 0.00202232 | 0.04615325 | *Rps19/Rpl30/Gapdh* | 3 |
| GO:0044364 | disruption of cells of other organism | 3/148 | 0.00202232 | 0.04615325 | *Rps19/Rpl30/Gapdh* | 3 |

**Table S5.** Primers and conditions used for real-time PCR.

| **Gene name** | **Primer sequence** | **Concentration used** | **Annealing temperature** |
|---|---|---|---|
| *Stat4* | F: GACCCTGAAGGCCGATTCTG | 200nM | 60$^{o}$C |
| | R: TGATTCCACTGAGACATGCTGG | 200nM | |
| *Actb* | F: AACGGCTCCGGCATGTGCAAAG | 200nM | 60$^{o}$C |
| | R: ATCACACCCTGGTGCCTAGGGCG | 200nM | |
| *18S* | F: TTCGAGGCCCTGTAATTGGA | 200nM | 60$^{o}$C |
| | R: GCAGCAACTTTAATATACGCTATTGG | 200nM | |

### Figures

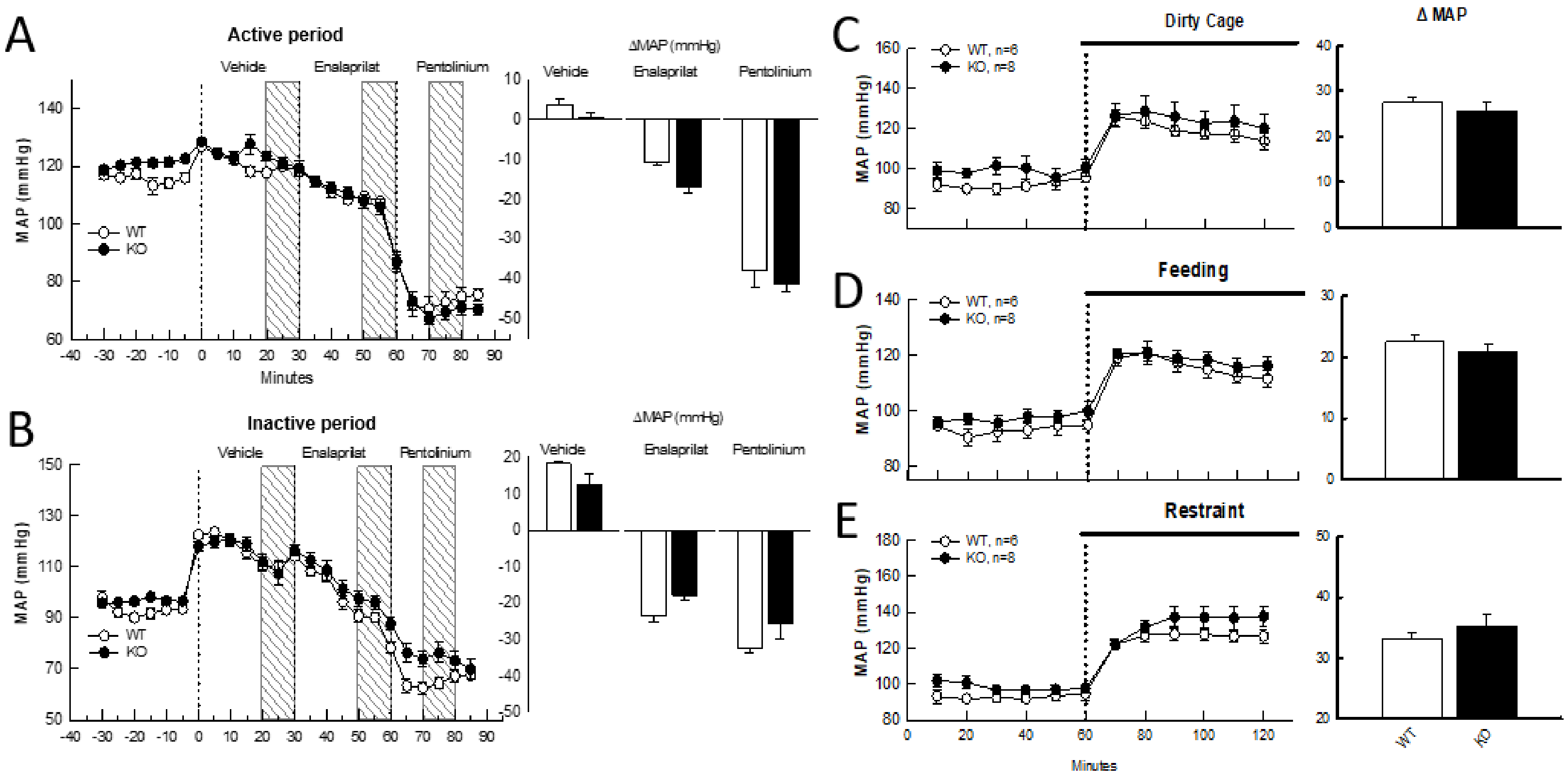


**Figure S1**. Changes in mean arterial pressure (MAP) in response to successive injections of vehicle (0.9% saline), enalaprilat and pentolinium during the **A,** active and **B**, inactive period in WT (n=6) and KO (n=8) mice. Dotted lines represent the time at which each i.p. injection was given. The hatched region shows the period which was used to determine the average change in MAP in response to vehicle, enalaprilat and pentolinium, as shown in the histograms. Changes in MAP in response to **C,** dirty cage-switch, **D**, feeding and **E**, restraint in WT (n=6) and KO (n=8) mice. All values are presented as mean ± SEM.

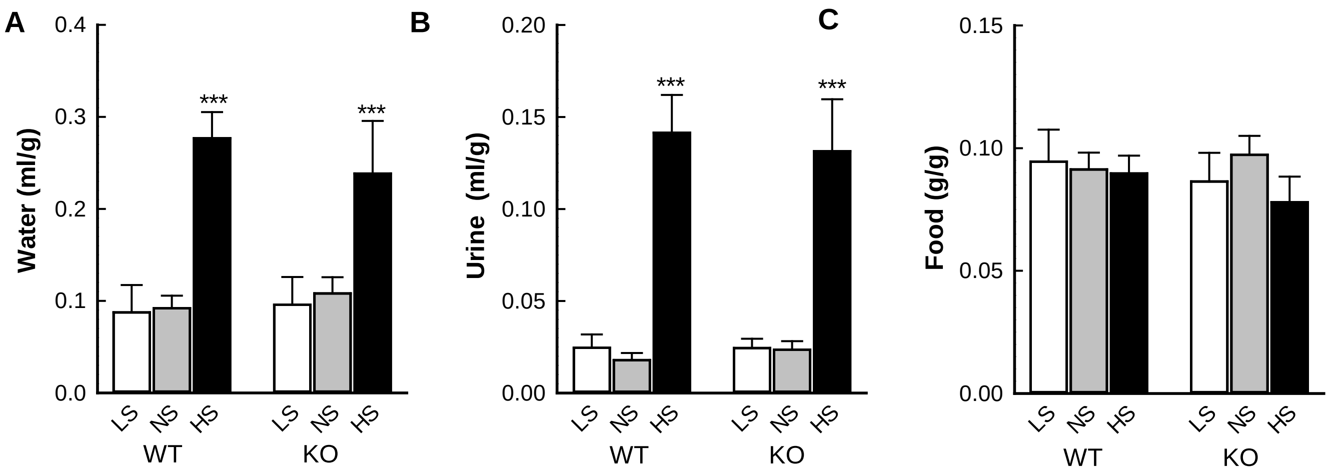


**Figure S2.** Average **A,** water consumption **B,** urine output and **C,** food intake in WT (n=6) and KO (n=6) mice in response to low (LS), normal (NS) and high (HS) salt diets. Bars represent average values ± SEM. Statistical analysis was conducted using split-plot analysis of variance. Comparisons are within strains ****P*<0.001 compared to NS.

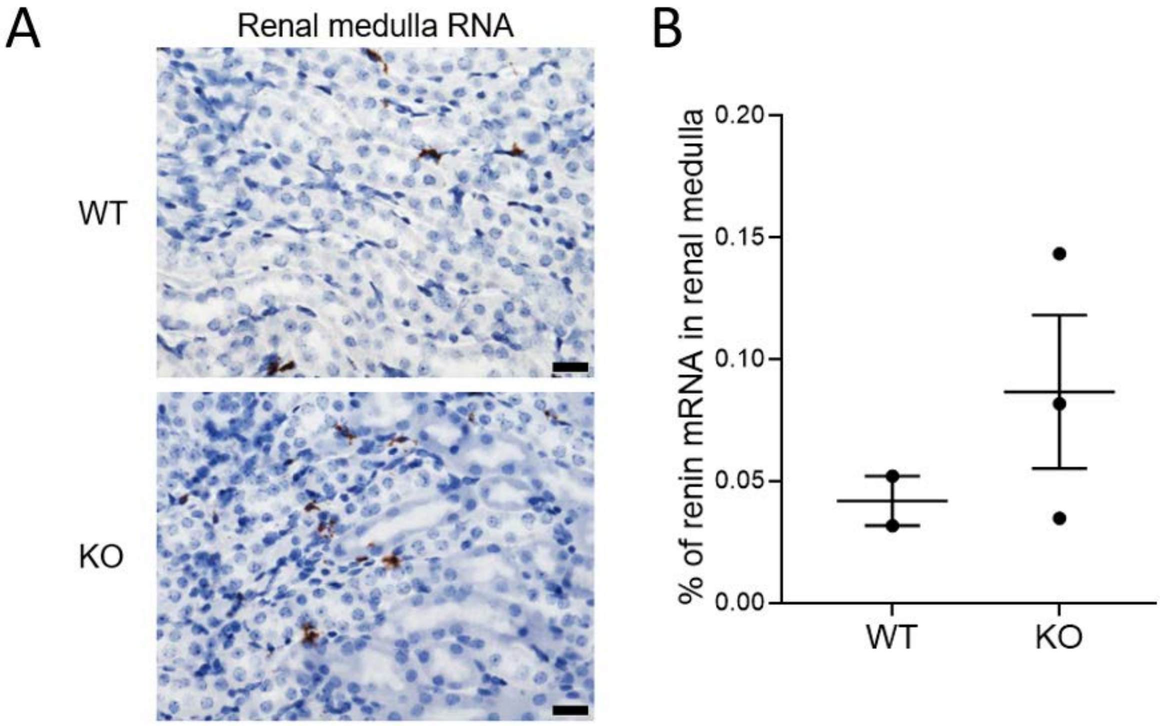


**Figure S3.** Percentage of renin mRNA in renal medulla of wild-type (WT) and miR-181a knockout 9KO) mice. This measurement was only available in 2 samples from WT mice, so statistical analyses were not performed. Scale bar= 20 µm.

A

| Parameters | WT | KO |
|---|---|---|
| Estimated Number of Cells | 3,361 | 3,338 |
| Mean Reads per Cell | 139,800 | 140,517 |
| Median Genes per Cell | 955 | 1,021 |
| Number of reads | 469,869,317 | 469,044,343 |
| Sequencing saturation | 96.2% | 95.8% |
| Fraction reads in cell | 89.1% | 90.2% |
| Total genes detected | 17,974 | 18,243 |
| Median UMI counts per cell | 2,214 | 2,344 |

B

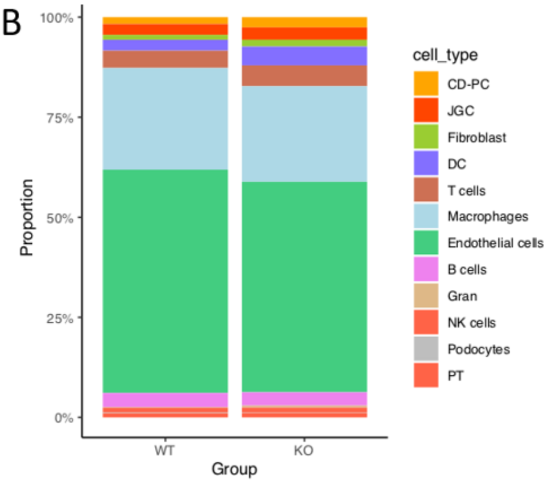


**Figure S4.** Summary of relevant single-cell RNA-sequencing data. A, Quality control data from single-cell RNA-sequencing data. B, distribution of cell types within all cells sequenced. Legend: WT, wild-type; KO, miR-181a knockout; Gran, granulocytes; PT, proximal tubules; NK cells, natural killer cells; DCs, dendritic cells tubule; JGC, juxtaglomerular cells; CD-PT, collecting duct principal cells.

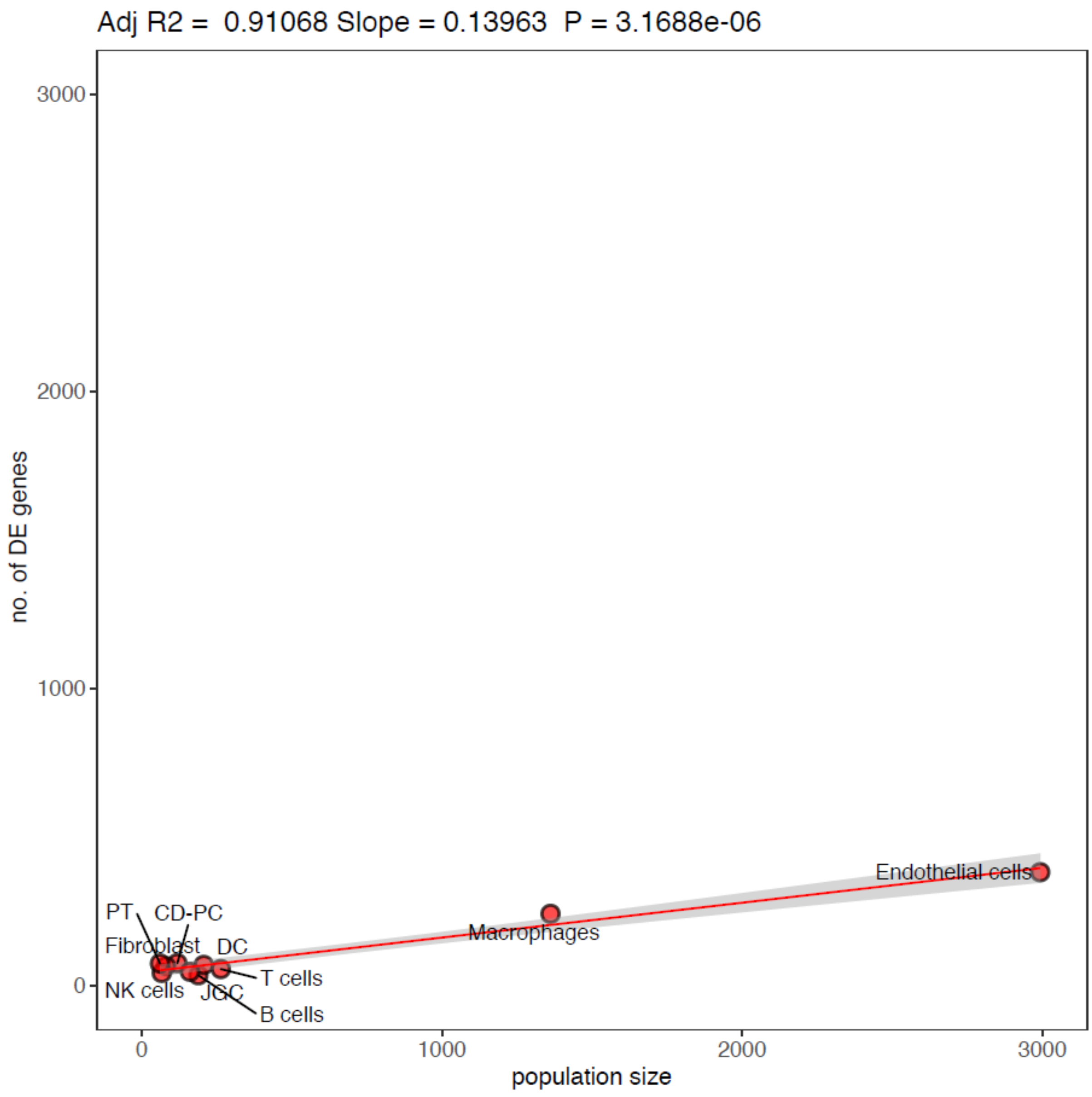


**Figure S5.** Number of differentially expressed genes according to cell population size.

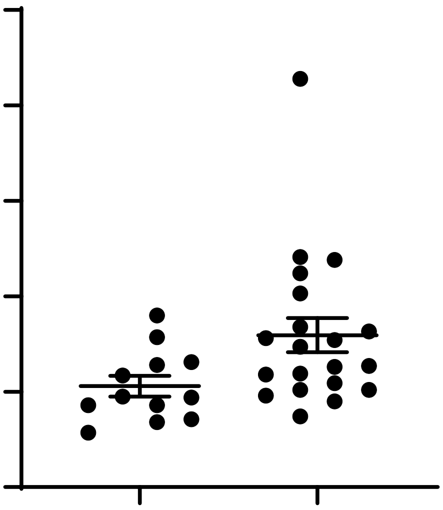

**Figure S6.** Validation of the expression of *Stat4* mRNA in whole kidney tissue using real-time PCR. WT mice (n=12) had lower levels of *Stat4* mRNA compared to miR-181a KO mice (n=20). t-test showing **$P$<0.01.
